# Diverse transcriptomic signatures across human tissues identify functional rare genetic variation

**DOI:** 10.1101/786053

**Authors:** Nicole M. Ferraro, Benjamin J. Strober, Jonah Einson, Xin Li, Francois Aguet, Alvaro N. Barbeira, Stephane E. Castel, Joe R. Davis, Austin T. Hilliard, Bence Kotis, YoSon Park, Alexandra J. Scott, Craig Smail, Emily K. Tsang, Kristin G. Ardlie, Themistocles L. Assimes, Ira Hall, Hae Kyung Im, GTEx Consortium, Tuuli Lappalainen, Pejman Mohammadi, Stephen B. Montgomery, Alexis Battle

## Abstract

Rare genetic variation is abundant in the human genome, yet identifying functional rare variants and their impact on traits remains challenging. Measuring aberrant gene expression has aided in identifying functional, large-effect rare variants. Here, we expand detection of genetically driven transcriptome abnormalities by evaluating and integrating gene expression, allele-specific expression, and alternative splicing from multi-tissue RNA-sequencing data. We demonstrate that each signal informs unique classes of rare variants. We further develop Watershed, a probabilistic model that integrates multiple genomic and transcriptomic signals to predict variant function. Assessing rare variants prioritized by Watershed in the UK Biobank and Million Veterans Program, we identify large effects across 34 traits, and 33 rare variant-trait combinations with both high Watershed scores and large trait effect sizes. Together, we provide a comprehensive analysis of the transcriptomic impact of rare variation and a framework to prioritize functional rare variants and assess their trait relevance.

**One-sentence summary:** Integrating expression, allelic expression and splicing across tissues identifies rare variants with relevance to traits.

## Background

Any given human genome contains tens of thousands of rare variants (*1, 2*) and rare variation contributes to both rare and common disease risk (*3, 4*). However, identifying high-impact rare variants, especially in the noncoding genome, remains difficult due to their low frequency and the lack of a regulatory genetic code. Previous work has shown that abnormal gene expression aids in identifying functional, large-effect rare variants (*5–9*). However, transcriptome sequencing provides diverse measurements beyond gene expression level including allele-specific expression and alternative splicing that have yet to be systematically evaluated and integrated into variant effect prediction (*10–12*). Furthermore, ongoing expansion of population-scale datasets with both whole genome and transcriptome data provides new opportunities to study the molecular impact of rarer variants.

With the substantial number of both whole genome and transcriptome samples (n=838) in the Genotype Tissue Expression (GTEx) project v8, we assess variants deeper into the frequency spectrum to analyze how novel and rare genetic variants contribute to outlier patterns in total expression (termed “expression”), allelic expression, and alternative splicing. We integrate these three transcriptomic signals across 49 tissues along with diverse genomic annotations to prioritize high impact rare variants, and assess their relationship to complex traits in the UK Biobank (*13*) and the Million Veterans Program (*14*).

## Results

### Detection of aberrant gene expression across multiple transcriptomic phenotypes

We quantified three transcriptional phenotypes for each gene to capture a wide range of functional effects caused by regulatory genetic variants. Briefly, to identify expression outliers (eOutliers), we corrected expression measurements for age, sex, genetic ancestry, hidden factors and the strongest cis-eQTL per gene (see Supplementary Methods, Fig S1) across the population and generated Z-scores to determine whether a gene in an individual has an extremely high or low expression (*15*). Splicing outliers (sOutliers) were detected using SPOT (SPlicing Outlier deTection), a novel approach which fits a Dirichlet-Multinomial distribution directly to counts of reads split across alternatively spliced exon-exon junctions for each gene. SPOT then identifies individuals that deviate significantly from the expectation based on this fitted distribution (see Supplementary Methods, Figs S2, S3). Finally, to identify genes with excessive allelic imbalance (aseOutliers) we used ANEVA-DOT (see Supplementary Methods, Fig S4, S5). This method uses estimates of genetic variation in dosage of each gene in a population to identify genes where an individual has a heterozygous variant with an unusually strong effect on gene regulation (*16*). Each of the three methods was applied across all GTEx samples; an individual is called an “outlier” for a method if their median value across measured tissues exceeds thresholds for a given gene (Fig 1A).

**Figure 1.**
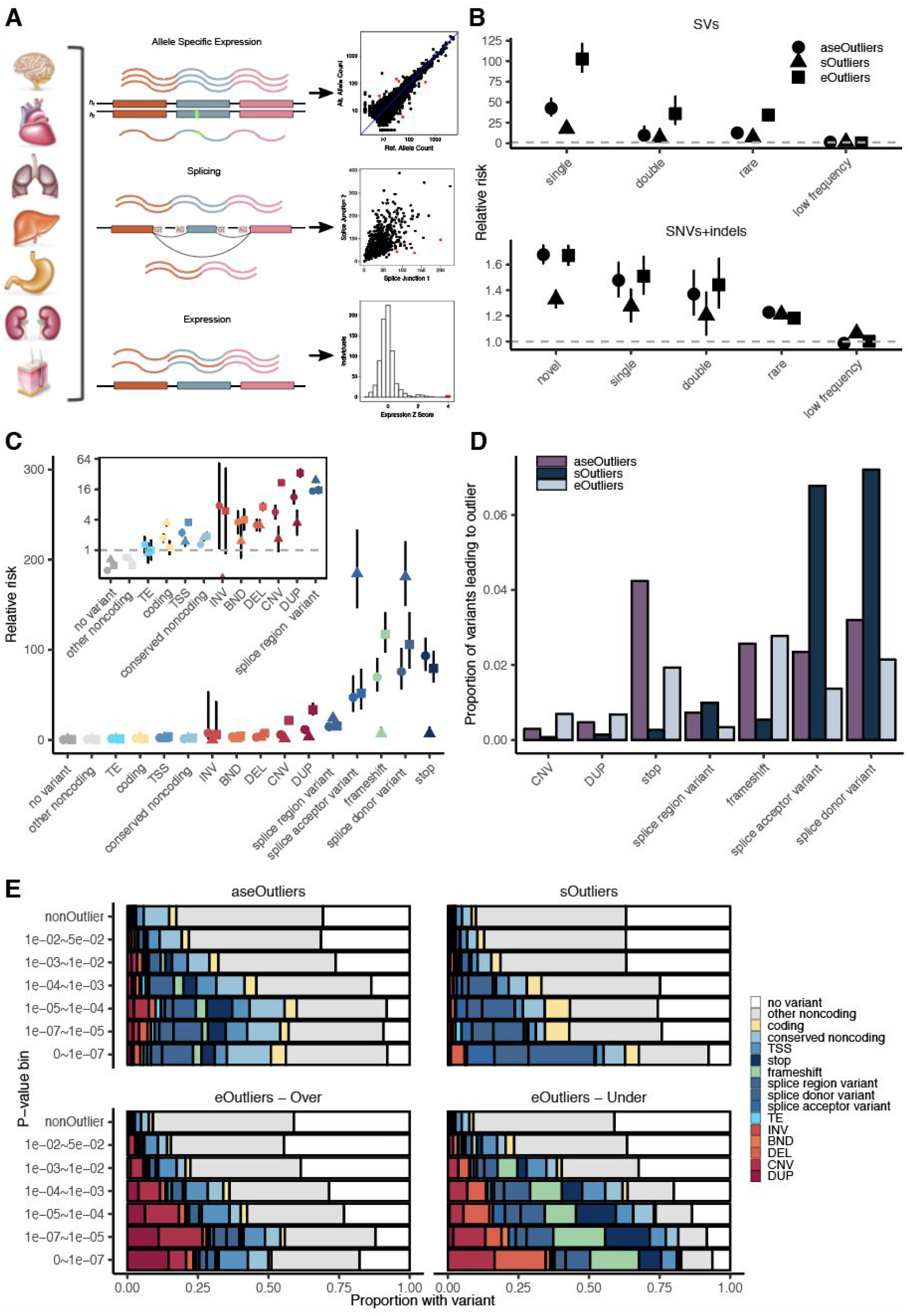
Enrichment of rare variants underlying aberrant expression, splicing, and allele specific expression. **(A)** RNA-sequencing data in 838 individuals was combined across 49 tissues and used to identify shared tissue expression, alternative splicing and ASE outliers. **(B)** Enrichment of novel (not in gnomAD), singleton, doubleton, rare (MAF < 1%) and low frequency (MAF 1-5%) variants across the allele frequency spectrum in a 10kb window around the outlier genes across all data types. Outliers were defined as those with values greater than 3 standard deviations from the mean (|MedZ| > 3), or equivalently, a median p-value < 0.0027. Bars represent the 95% confidence interval. **(C)** Assigning each outlier its most consequential nearby rare variant, enrichments for different categories of rare variants for each outlier type. **(D)** The proportion of rare variants in a given category that lead to an outlier at a p-value threshold of 0.0027 across types. **(E)** Proportion of outliers at a given threshold that have a nearby rare variant in the given category. |Median Z-scores| are converted to p-values for expression outliers. TSS = transcription start site, TE = transposable element, INV = inversion, BND = breakend, DEL = deletion, CNV = copy number variation, DUP = duplication

### Aberrant expression, ASE and splicing genes are enriched for distinct rare variants

We observed that outlier individuals for any of the three transcriptomic phenotypes are significantly more likely to carry a rare variant (MAF < 1%) in the gene body or +/− 10kb than individuals without outliers, assessed among 714 individuals with European ancestry. While we saw enrichments for all variant classes and outliers up to a 1% allele frequency, the strongest enrichments are for variants of the lowest frequencies. Rare structural variants seen only once within GTEx are most strongly enriched, with 103-fold (eOutliers), 42.6 (aseOutliers), and 17.6-fold (sOutliers) enrichments respectively over non-outlier individuals (Fig 1B). Overall, structural variants and ultra-rare variants exhibit disproportionately significant impact on the transcriptome.

For the set of individuals and genes tested by all methods, 35 outliers were found by all three methods, all but one of which have a nearby rare variant with half annotated as splice variants. For outliers detected across molecular phenotypes, the greatest sharing occurs between aseOutliers and eOutliers. We found that aseOutliers tend to show an impact on expression even when it is too modest to reach our threshold for detecting eOutliers (Wilcoxon rank-sum p-value < 2.2 × 10^−16^). Furthermore, aseOutliers with modest expression changes, 1 < |median Z| < 3, showed stronger enrichment for nearby rare variants than those without an expression change (Fig S6), highlighting a benefit of combining these phenotypes to discover diverse rare variant effects.

We found that different types of genetic variants contribute to outliers for the three molecular phenotypes. Splice donor variants are strongly enriched nearby all outlier types. For aseOutliers, the strongest enrichments occur for stop-gained variants at 93.2-fold, while sOutliers show strongest nearby enrichments for splice acceptor variants at 185-fold and eOutliers for frameshift variants at 117-fold (Fig 1C). The largest differences among data types were copy number variations (CNVs) and duplications that were enriched within eOutliers, and splice acceptor variants that were enriched within sOutliers (Fig S7). All outlier types were significantly depleted for having no rare variants in the 10kb window around the gene and also depleted for non-coding rare variants that do not fall in any other annotated category (Fig 1C).

For all phenotypes, the proportion of outliers with a nearby rare variant of any category increases with threshold stringency. We split eOutliers by over- and under-expression. At the strictest threshold, of 336 over-eOutliers 82% have a nearby rare variant, of which 62% have an additional annotation (from those in Fig 1C) and of 175 under-eOutliers, 94% have a nearby rare variant, 88% of which are annotated. Of 584 aseOutliers, 92% have a nearby rare variant, of which 56% are annotated as at least conserved non-coding. For sOutliers, of 644, 92% have a nearby rare variant of which 73% are annotated (Fig 1D). This analysis provides additional insight into expectations for causal rare variant types when an outlier effect of a specific magnitude is observed in an individual.

While the strongest outliers indicate the presence of rare genetic variants well over 75% of the time, a large proportion of rare genetic variants do not manifest as large molecular aberrations, even for the most predictive classes of variants. For example, for any given rare variant occurring within 10kb of a gene, for eOutliers, rare frameshift variants are most strongly enriched, but at just a 2.8% absolute frequency. Rare splice donor variants lead to an aseOutlier in 3.2% of instances and rare stops are associated with aseOutliers in 4.2% of instances. For sOutliers, 7.2% of rare splice donor variants lead to a sOutlier, as will 6.8% of rare splice acceptor variants (Fig 1E). When randomly selecting an equal number of gene-individuals as outliers over 1000 iterations, percentages greater than or equal to these were never observed, reflecting true enrichment for each category (Fig S7). Overall, the low frequency with which rare variants of these classes lead to large transcriptome changes highlights the utility of incorporating functional data in variant interpretation even for some variant classes already used in clinical interpretation. While subtle or context-specific transcriptomic effects may sometimes escape detection due to noise in RNA-sequencing data, tissue selection, or methodological choices, many annotated likely gene disrupting variants may actually not be functional (*17*). In those situations, incorporating functional data in variant interpretation will help to pinpoint variants with true functional effects. We return to this point in subsequent sections, developing methods to identify variants with much greater absolute risk of functional effects.

### Genomic position of rare variants predicts impact on expression

While we primarily assessed rare variants that occur either within an outlier gene, or in a 10kb surrounding window, gene regulation can occur at greater distances (*18, 19*). As we observed the strongest enrichments for the lowest frequency variants, we intersected singleton variants (SVs appearing only once in GTEx, and SNVs/indels that do not appear in gnomAD) with 200kb length windows, exclusive of other windows, upstream from outlier genes and compared their frequency in outlier vs non-outlier individuals. SNV enrichments dropped off quickly at greater distances from the gene, though remained weakly enriched for eOutliers out to 200kb. The same is true for rare indels, with enrichment at 200kb only for sOutliers. However, singleton SVs remained enriched at longer distances, and are enriched at 2.33-fold as far as 800kb-1Mb (Fig 2A). Thus, transcriptome data can help detect distal cis-regulatory effects that would often be difficult to annotate and link to target genes using only genetic data.

**Figure 2.**
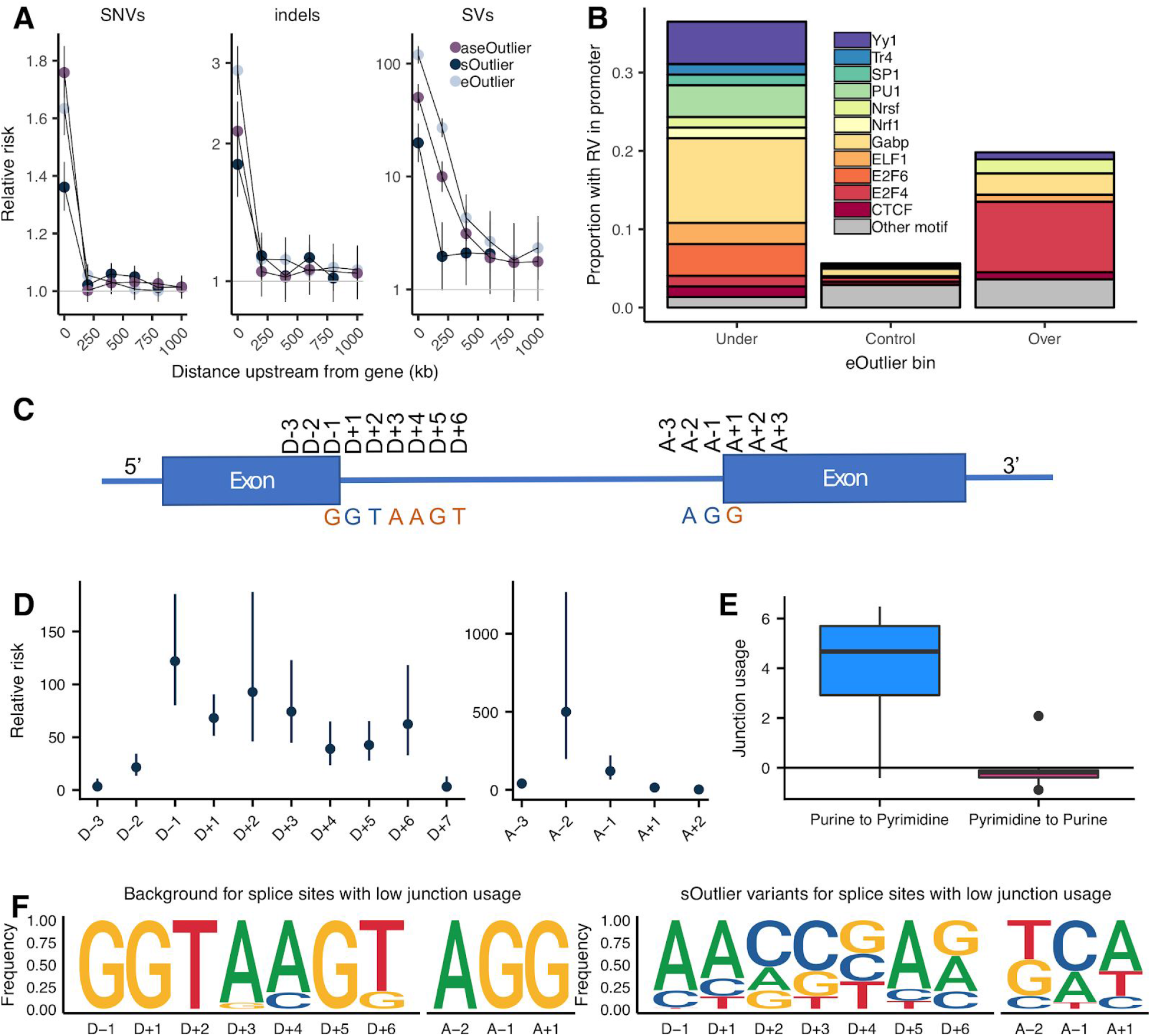
Rare variant enrichments in specific genomic positions. **(A)** Enrichment of singleton SNVs, indels, and SVs at varying distances upstream of outlier genes (bins exclusive) across data types. **(B)** Enrichment of TSS rare variants in promoter motifs within 1000bp of eOutliers. Under and over bins are defined using a median Z-score threshold of 3, and controls are all individuals with a median Z-score less than 3 for the same set of outlier genes. **(C)** Graphic summarizing positional nomenclature relative to observed donor and acceptor splice sites. **(D)** Relative risk (y-axis) of a sOutlier (median LeafCutter cluster p-value < 1 × 10^−5^) rare variant being located at a specific position relative to the splice site (x-axis) compared to non-outlier rare variants. Relative risk calculation done separately for donor and acceptor splice sites. **(E)** Junction usage of a splice site is the natural log of the fraction of reads in a LeafCutter cluster mapping to the splice site of interest in sOutlier (median LeafCutter cluster p-value < 1 × 10^−5^) samples relative to the fraction in non-outliers samples aggregated across tissues by taking the median (see Supplementary Methods). Junction usage (y-axis) of the closest splice sites to rare variants that lie within a polypyrimidine tract ([A-5, A-35]) binned by the type of variant (x-axis). **(F)** Independent position weight matrices showing mutation spectra of sOutlier (median LeafCutter cluster p-value < 1 × 10^−5^) rare variants at positions relative to splice sites with negative junction usage (ie. splice sites used less in outlier individuals than in non-outliers).

Rare variants in promoter regions have been previously linked to outlier expression (*6, 15*). To extend these observations and assess the types of transcription factor binding sites that could lead to outliers, we tested enrichment of rare TSS variants in several specific TF motifs nearby under- and over-eOutliers. For under-eOutliers, we saw an enrichment of variants in *GABP* that activates genes that control cell cycle, differentiation, and other critical functions (*20*). For over-eOutliers, we saw an enrichment of rare variants intersecting the *E2F4* motif that has been reported as a transcriptional repressor (*21*). In both under and over-eOutliers, we saw rare variants in *YY1* that can act as both an activator or repressor, depending on context (*22*) and has been associated with *GABP* in co-regulatory networks (*23*) (Fig 2B). Thus, these naturally occurring rare variant perturbations can provide information about how specific TFs up- and down-regulate their target genes.

We observed that some rare variants can impact multiple genes potentially yielding oligogenic effects. We found a strong enrichment for eOutliers, and to a lesser degree, aseOutliers, occurring in the same individual within close proximity. As expected, we do not see a similar enrichment for nearby sOutlier pairs, which are less subject to co-regulation. Within a 100kb window, neighboring eOutlier genes are 70 times more frequent than expected by chance. These outlier gene pairs are enriched for rare CNVs, duplications and TSS variants nearby one or both genes, as compared to individuals who have outlier expression for only one of the genes (Fig S8), providing insight into transcriptional expectations for genes in the vicinity of these types of rare variants.

Previous studies have shown rare variants disrupting splice sites result in outlier alternative splicing patterns (*24, 25*). We used sOutlier calls made for each LeafCutter cluster (see Supplementary methods), providing greater resolution than aggregating by gene, to more precisely assess enrichment of splicing-related variants. As expected, we observed extreme enrichment of rare variants near splice sites in sOutliers. A sOutlier is 333 times more likely than a non-outlier to harbor a rare variant within a two base pair window around a splice site (Fig S9A; see Supplementary methods), with signal decaying at greater distances but still modestly enriched (relative risk 7.43) up to 100 bp away. To get base pair resolution enrichments, we computed the relative risk of sOutlier rare variants located at specific positions relative to observed donor and acceptor splice sites (see Supplementary methods). Ten positions near the splice site show significant enrichment for harboring a sOutlier variant (Fig 2C,D), corresponding precisely to positions that have also been shown to be intolerant to mutations, due to their conserved role in splicing (we will refer to these positions as the splicing consensus sequence) (*25*). Among the most enriched positions within the splicing consensus sequence were the four essential splice site positions (D+1, D+2, A-2, A-1) (*26*), showing an average relative risk of 195.

sOutliers further captured the transcriptional consequences both for variants that disrupt a reference splicing consensus sequence, as well as for variants that created a new splicing consensus sequence that was not previously present. Individuals with sOutlier variants where the rare allele deviates away from the splicing consensus sequence showed decreased junction usage of the splice site near the variant, whereas individuals with variants where the rare allele creates a splicing consensus sequence showed increased junction usage of the splice site near the variant relative to non-outliers (see Supplementary methods; Fig 2F, Fig S9B, Fig S10). We saw a related enrichment pattern after separating annotated and novel (unannotated) splice sites. (Fig S11). sOutliers were also enriched for rare variants positioned within the Polypyrimidine Tract (PPT). The PPT is a highly conserved, pyrimidine rich region, approximately 5 to 35 base pairs upstream from acceptor splice sites (*27*). A rare variant is 6.25 times more likely to be located in the PPT near an sOutlier relative to a non-outlier. sOutliers with a rare variant that changes a position in the PPT from a pyrimidine to a purine (ie. disrupting an existing PPT) show decreased junction usage of the splice site near the variant, whereas the inverse is true for variants that change a position in the PPT from a purine to pyrimidine (Fig 2E, Fig S12).

### Rare variants in tissue-specific regulatory regions can lead to tissue-specific aberrant expression

While outliers detected across most tissues or samples from the same individual offer improved power to detect rare variant effects, we are also now able to evaluate rare variants based on outliers detected in individual tissues. First, we performed replication analysis across all individuals with data available for the three methods to evaluate the degree to which outlier status detected in one tissue of an individual is replicated in other tissues. Among all individual-gene outliers across all methods in a discovery tissue, we calculated the percentage of times the same individual-gene pair was detected as an outlier in a test tissue, limiting to tissues where both genes are expressed. We then aggregated this calculation across all individuals and genes (Figure 3A, S14). On average, we find that the eOutlier, aseOutlier, and sOutlier status in a discovery tissue is detected in a test tissue 5.1%, 10.7%, and 8.7% of the time, respectively. This is consistent with other findings that measurements of ASE better replicate across tissues (The GTEx Consortium 2019, in submission) and has implications for the use of functional data from easily accessible tissues to understand disease states.

**Figure 3.**
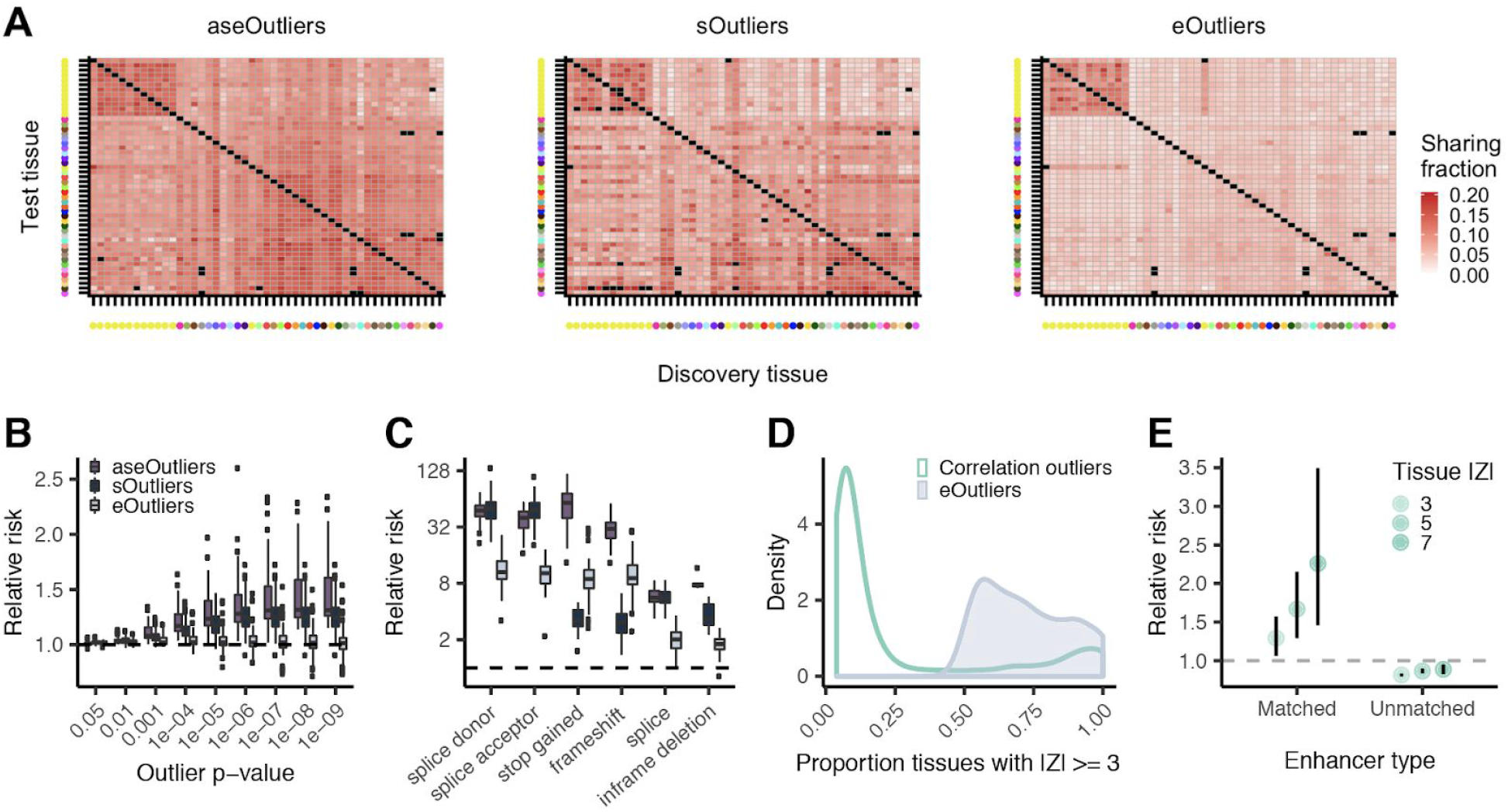
Single-tissue outlier enrichments and replication. **(A)** Outlier replication across tissues for each outlier type. Black indicates untested pairs. Tissue color legend in Fig S14. **(B)** Relative risk point estimate for nearby rare SNVs for outliers across all tissues individually. **(C)** Relative risk enrichments for likely gene disrupting nearby single-tissue outliers at a threshold of |Z| > 4 (p < 0.000063), with one point per tissue. **(D)** Distribution of number of tissues with aberrant expression underlying expression outliers defined via median Z-score (MEDZ) or Mahalanobis distance p-value (Correlation). **(E)** Enrichment of correlation outliers driven by a single tissue for rare variants in enhancers annotated to that tissue within a 500kb window of the outlier gene. Unmatched are defined as all tissue-specific enhancer regions, regardless of outlier tissue.

We next evaluated the ability of single-tissue outliers from each method to prioritize rare variants near outlier genes. Single-tissue aseOutliers were most enriched for nearby rare variants, followed by sOutliers than eOutliers, across all outlier cutoff thresholds (Figure 3B, Fig S14). We also observed enrichment of variants likely triggering nonsense mediated decay (NMD) among single-tissue aseOutliers, sOutliers, and eOutliers, among other predicted functional annotations, indicating that the most likely deleterious variants can still have a measurable effect in a single tissue (Figure 3C). Additionally, we found that sOutliers, even when restricted to evaluation of a single tissue, show extreme enrichment for rare variants in the splicing consensus sequence and the PPT (Fig S13). These results indicate that multi-tissue analysis substantially increases the detection of rare genetic effects, but transcriptome data from a single tissue does also inform on rare functional variants.

We did not see rare variant enrichments for eOutliers in single tissues until the most extreme thresholds, except for rare structural variants which are still enriched at comparable thresholds to multi-tissue outliers, though to a lesser degree (Fig S15). Therefore, we applied a different approach to discover tissue-specific expression effects by calling outliers based on their deviation from the expected covariance of expression across tissues on a gene by gene basis (see Supplementary Methods). A similar approach has been implemented to identify functional rare variants based on the correlation of expression among genes in a single tissue (*6*). Here we leveraged the breadth of tissue data available and used normal patterns of correlation across tissues to detect outliers that deviate in subsets of tissues in unexpected patterns to help identify tissue-specific functional rare variants. We found outliers identified via this approach were often driven by expression changes in one or a few tissues as compared to |median Z| outliers, which by definition, require changes in at least half of tested tissues (Fig 3C). These outliers were enriched for nearby rare variants in a 10kb window around the gene (Fig S16C). However, tissue-specific correlation outliers were also enriched for rare variants in enhancers that are active in the tissue driving the outlier effect, as determined by single-tissue Z-score, within a 500kb window around the gene, as compared to any enhancer region present in any tissue (Fig 3D). These tissue-specific outliers were depleted for rare variation in enhancers annotated to other tissues, highlighting the importance of considering tissue-specific regulatory annotations when assessing tissue-specific expression outliers.

### Prioritizing rare variants by integrating genomic annotations with diverse personal transcriptomic signals

To incorporate diverse transcriptome signals into a method to prioritize rare variants, we developed Watershed, an unsupervised probabilistic graphical model that integrates information from genomic annotations of a personal genome with multiple signals from a matched personal transcriptome. Watershed provides scores that can be used for personal genome interpretation, describing the posterior probability given both WGS and RNA-seq signals that a variant has a functional effect on each transcriptomic signal (Fig 4A). The Watershed model can be adapted to any available collection of molecular phenotypes, whether different assays, different tissues, or different derived signals. Further, Watershed automatically learns Markov random field (MRF) edge weights reflecting the strength of the relationship between the different phenotypes included that together allow the model to optimally predict functional effects.

**Figure 4.**
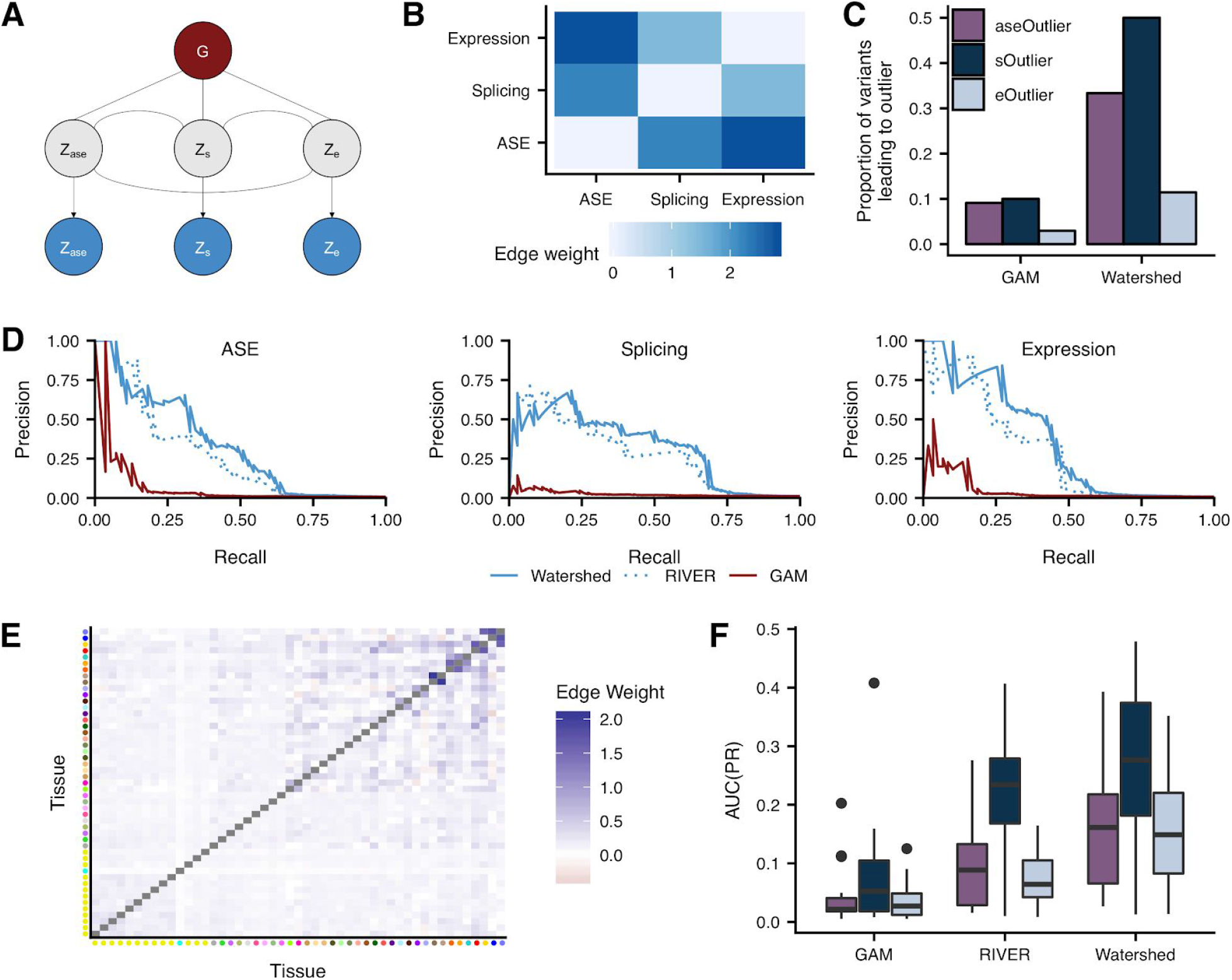
Prioritizing functional rare variants with Watershed. **(A)** Graphic summarizing plate notation for Watershed model when it is applied to three median outlier signals (ASE, Splicing, and Expression). **(B)** Learned Watershed edge parameters (weights) between pairs of outlier signals after training Watershed on three median outlier signals. **(C)** The proportion of rare variants, with Watershed posterior probability greater than 0.9 (right) and with genomic annotation model (GAM) probability greater than a threshold set to match the number of Watershed variants for each outlier signal (left), that lead to an outlier at a median p-value threshold of 0.0027 across three outlier signals (colors). Watershed and GAM models evaluated on held-out pairs of individuals. **(D)** Precision-recall curves comparing performance of Watershed, RIVER, and GAM (colors) using held out pairs of individuals for three median outlier signals. **(E)** Learned tissue-specific Watershed edge parameters (weights) between pairs of tissue outlier signals after training tissue-specific Watershed on eOutliers across single tissues. Tissue color to tissue name mapping can be found in Supplementary Figure S14C. **(F)** Area under precision recall curves (AUC(PR); y-axis) in a single tissue between GAM, RIVER, and tissue-specific Watershed (x-axis) when applied to outliers across single tissues in all three outlier signals (colors). Precision recall curves in each tissue generated using held out pairs of individuals.

We first applied Watershed to the GTEx v8 data using the three outlier signals examined here: ASE, splicing, and expression (Fig 4A; see Supplementary Methods), where each was first aggregated by taking the median across tissues for the corresponding individual. In agreement with existing evidence of similarity between outlier signals (Fig S6), the learned Watershed edge parameters were strongest between ASE and expression but strictly positive for all pairs of outlier signals (ie. each outlier signal is informative of all other signals; Fig 4B). To evaluate our model we used held out individuals (see Supplementary Methods). Watershed greatly outperforms methods based on genome sequence alone (our Genomic Annotation Model (GAM) and CADD; Fig S17 and Fig 4C) (*28, 29*). Explicitly modeling the relationship between different molecular phenotypes also offers a performance gain (Watershed compared to RIVER; Fig 4D, Fig S18). We have shown that even the most predictive genomic annotations only result in a transcriptomic outlier less than 10% of the time (Fig 1E and Fig 4C). However, integrating transcriptomic signals with genomic annotations via Watershed (at a posterior threshold of 0.9) detected SNVs that result in aseOutliers, sOutliers, and eOutliers with much greater frequency, 33.3, 50.0, 11.4% of the time, respectively (Fig 4C, Fig S19).

We further extended the Watershed framework to provide tissue-specific scores to rare variants. In this version, Watershed prioritizes variants based on their predicted impact on transcription in each individual tissue, while sharing evidence between related tissues. Specifically, we trained three independent tissue-specific Watershed models (one for ASE, splicing, and expression) where each model considers effects in all tissues, giving 49 phenotypes in each model (See Supplementary methods, Fig S18, Fig S20). In this context, MRF edge parameters can be interpreted as the strength that functional variant effects are shared between pairs of tissues. Edge parameters from each of the three tissue-specific Watershed models resemble known patterns of tissue similarity (Fig 4E, Fig S21). Further, using held out individuals, the tissue-specific Watershed model outperforms a comparable model where each tissue is treated completely independently (Fig 4F, Fig S22). Tissue-specific Watershed also performs better than a comparable model trained using single median outlier statistics (Fig S23). Critically, Watershed models trained on a single tissue independently do still outperform methods based only on genome sequence annotations, supporting the benefit of collecting even a single RNA-seq sample to improve personal genome interpretation. However, when available, multiple samples, tissues, or assays can be incorporated to further improve performance and provide tissue-(or assay-) specific predictions.

### Aberrant expression informs rare variant trait associations

Overall, we found that a given individual has a median of 13 aseOutliers, 19 sOutliers, and 11 eOutliers after filtering for those with a nearby rare variant. When filtering by moderate Watershed posterior probability (>0.5) of impacting ASE, splicing, or expression, an individual has a median of 2 rare variants predicted to impact ASE, 5 predicted to impact splicing and 8 predicted to impact expression (Fig 5A). In order to assess whether the identified rare functional variants from GTEx associate with traits, we intersected this set with variants present in the UK Biobank (UKBB) (*13*). We focused on a subset of 34 traits for which GWAS association had evidence of colocalizations with e/sQTLs (Table S2). Genes with a rare variant trait association are strongly enriched for their eQTLs colocalizing with GWAS signals for the same trait (The GTEx Consortium 2019, in submission) indicating that QTL evidence can be used to guide rare variant analysis. Rare variants nearby outliers that appear in GTEx and in UKBB had larger trait association effect sizes than background rare GTEx variants near the same set of genes across tested traits (p=3.51 × 10^−9^), with a shift in median effect size percentile from 46% to 53%. Additionally, outlier variants that fell in or nearby genes with eQTL or sQTL colocalization had even larger effect sizes (median effect size percentile 88%) than non-outlier variants (p=1.93 × 10^−5^) or outlier variants falling near any gene (p=4.88 × 10^−5^) not matched to a colocalizing trait (Fig 5B).

**Figure 5.**
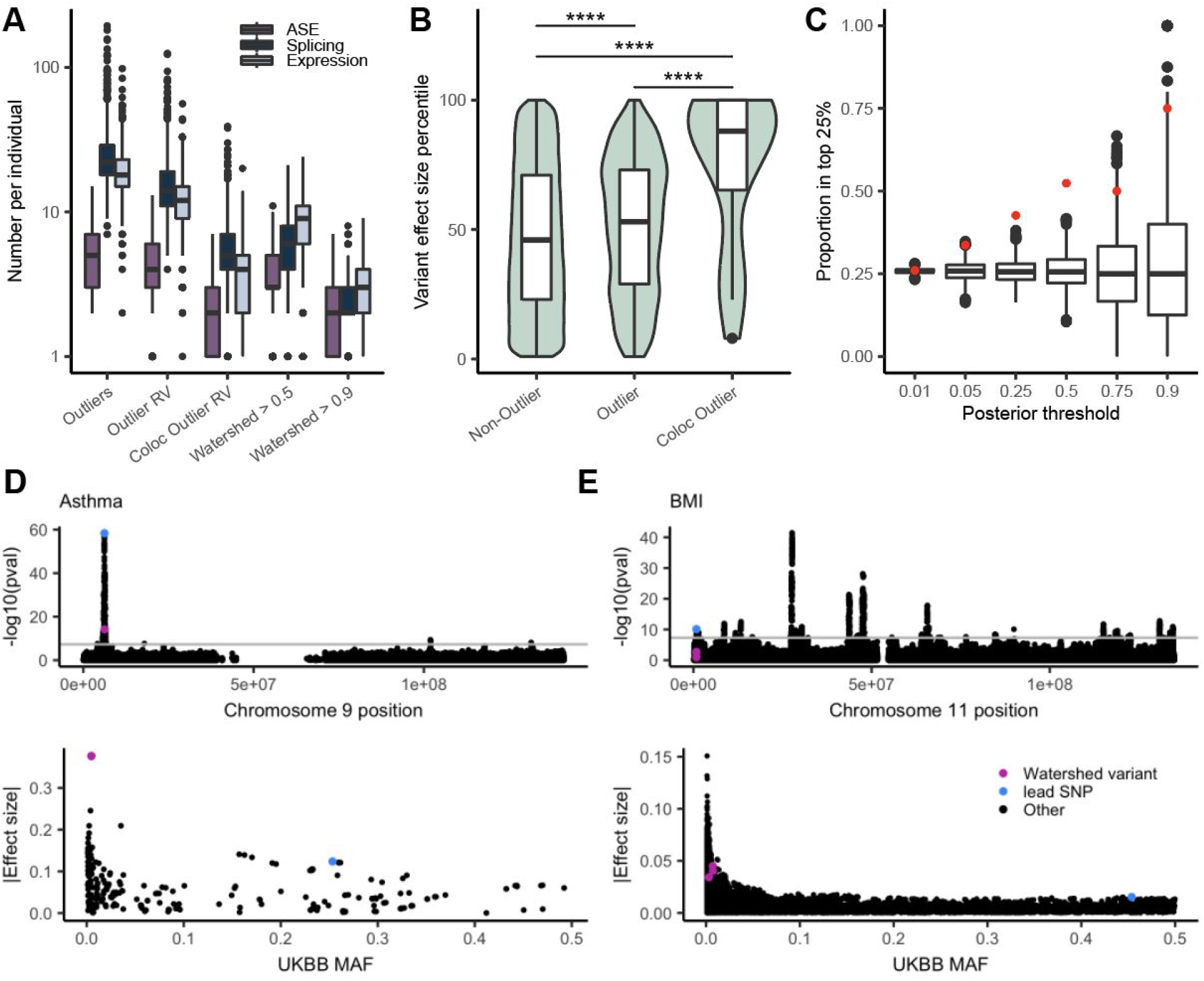
Trait associations for rare variants underlying outlier genes. **(A)** Distribution of the number of outliers, outliers with a nearby rare variant, outliers with a nearby rare variant in colocalizing genes, and variants with high Watershed score per data type. **(B)** The distribution of effect sizes, transformed to a percentile, for the set of GTEx rare variants that appear in UKBB and are not outlier variants, those that are outlier variants, and those outlier variants that fall in colocalizing genes for the matched trait across 34 traits. Percentiles are calculated on the set of rare GTEx variants that overlap UKBB. The set of genes is restricted to those that have at least one outlier individual in any data type, and a nearby variant included in the test set (4,787 variants and 1,323 genes). p-values are calculated from a one-sided Wilcoxon rank-sum test. **(C)** The proportion of variants filtered by Watershed posterior that fall in the top 25% of effect sizes for a colocalized trait (red), and the proportion of randomly selected variants of an equal number that also fall in these regions over 1000 iterations (black). **(D)** Manhattan plot (top) across chromosome 9 for asthma in the UKBB, filtered for non-low confidence variants, with the high-Watershed variant, rs146597587, in pink and the lead colocalized variant, rs3939286, in blue. This variant’s effect size rank was similarly high for both self-reported and diagnosed asthma, but we show the summary statistics for asthma diagnosis here. The UKBB MAF vs absolute value of the effect size for all variants within 10kb of the Watershed variant (bottom). **(E)** Manhattan plot across chromosome 11 for BMI in the UKBB, filtered to remove low confidence variants, with the high-Watershed variants, rs35835984, rs184629044, and rs192797357 in pink and the lead colocalized variant, rs6591, in blue. The UKBB MAF vs absolute value of the effect size for all variants within 15kb of the Watershed variant to capture the lead colocalized variant (bottom).

While most variants tested have low Watershed posterior probabilities of impacting the transcriptome (Fig S24A; termed Watershed scores), we hypothesized that filtering for those variants that do have high scores would yield variants in the upper end of the effect size distribution for a given trait. For each variant tested in UKBB, we took the maximum Watershed score per variant and compared this to a solely genomic annotation defined metric, CADD (*28, 29*). We find that Watershed scores were a better predictor of variant effect size than CADD scores for the same set of rare variants (Table 1). Furthermore, across different score thresholds, we found that the proportion of variants falling in the top 25% of rare variant effect sizes in colocalized regions exceeded the proportion expected by chance at most thresholds (Fig 5D). While filtering by CADD score did return some high effect size variants, this proportion declines at the highest thresholds (Fig S24D). Furthermore, there was very little overlap between variants with high Watershed scores and high CADD variants (Fig S24D), with CADD variants more likely to occur in coding regions, and Watershed variants more frequent in non-coding regions (Fig S24D). Thus, the approaches largely identify complementary sets of variants for these traits.

**Table 1.** Watershed and CADD as predictors of variant effect size percentile. Coefficient estimate and 95% confidence interval from separate linear models with variant effect size percentile as the response and CADD or Watershed score (scaled to have a mean of 0 and sd of 1 so values are of comparable range) as the predictor for all tested variants in co-localized regions (n=5,277).

| Predictor | Beta | p-value | 95% confidence interval |
| --- | --- | --- | --- |
| Watershed score | 1.61 | $2.12 \times 10^{-6}$ | 0.95 - 2.27 |
| CADD score | 0.77 | $2.41 \times 10^{-2}$ | 0.10 - 1.43 |

We identified 33 rare GTEx variant-trait combinations where the variant has a Watershed score > 0.5 and falls in the top 25% of variants by effect size for the given trait (Table S3). We highlight two such examples, for asthma and BMI (Fig 5E-F), showing that while rare variants usually do not have the frequency to obtain genome-wide significant p-values, when we prioritize by the probability of impacting expression, we can identify those that have greater estimated effect sizes on the trait. In the case of rs146597587 in asthma, the effect size in UKBB is 3 times greater than the lead colocalized variant. This variant, rs146597587, is a high-confidence loss of function splice acceptor with an overall gnomAD allele frequency of 0.0019, and falls within the gene IL33, whose expression has been previously implicated in asthma (*30, 31*). For BMI, we find three rare variants with high effect sizes, rs35835984, rs184629044, and rs192797357, which each have between a 2-3 times greater effect size on the trait than the lead colocalized variant. These variants have a high predicted probability of impacting the expression of two colocalized genes, *CD151* and *POLR2L*. For one additional high Watershed variant, rs564796245, for *TTC38,* which had a high effect size for self-reported high cholesterol in the UKBB, we were able to test this variant against four related blood lipids traits in the Million Veterans Program (*32*). We found that for these traits, which include HDL, LDL, total cholesterol, and triglycerides, among rare (gnomAD AF < 0.1%) variants within a 250kb window of rs564796245, this variant is in the top 10% of variants by effect size, and for HDL specifically, it is in the top 1% (Fig S25). Given that this variant is rare in all cohorts, and that the traits are not exact matches, seeing consistent higher effect sizes across these related traits in a separate cohort provides additional evidence for this variant’s association with cholesterol.

Only four of the variants tested in UKBB have Watershed posterior probabilities > 0.9 for colocalized genes, but of those, three show high effect sizes for the relevant trait (Table S3). We are limited in the traits that we can assess here, and by the small overlap in rare variants observed between the datasets as few population-scale phenotyped and genome sequenced cohort are yet available. As datasets including genome sequencing, functional molecular data and trait and disease phenotype data grow, application of Watershed and other integrative methods would likely prove fruitful in prioritizing rare variants impacting traits, especially those in the lowest ends of the allele frequency spectrum.

## Discussion

Rare variants are abundant in human genomes yet have remained difficult to systematically study. With over 800 genomes matched with transcriptomes across 49 human tissues available in GTEx v8 data, we have shown that rare variation, including singleton variants, underlie extreme changes in the transcriptome of carrier individuals. To better capture the diversity of transcriptome aberrations, we applied ANEVA-DOT to test aberrant allele-specific expression, and presented SPOT, a novel approach for discovering aberrant alternative splicing. Together, we applied these complementary approaches along with multi-tissue expression outlier detection for each GTEx individual, to assess rare variant impact on the transcriptome. We found that the genomic context of each rare variant is predictive of the nature and magnitude of the observed change in expression, such as changes to splice junction usage or tissue-specific effects.

For each transcriptomic signal, we observed rare variant enrichment that increased with the extremity of the transcriptional aberration. Observed enrichments were strongest for ultra-rare variants and reflected the specific nature of each signal. Among eOutliers, we observed strong enrichment of structural variants including those that influenced multiple genes and across large distances. sOutliers were particularly enriched for variants that disrupted or created the splicing consensus sequence or the polypyrimidine tract. AseOutliers and sOutliers were particularly powerful signals of rare variant effects, with enrichment evident even with a single tissue, which has implications for studies where it is only possible to collect data from one or a few tissues. Furthermore, using a novel approach for assessing tissue-specific eOutliers, we observed rare variants enriched in tissue-matched enhancers. Evaluation of rare variants in distant regulatory elements remains an important challenge for further work, including incorporation of additional measurements of chromatin state and expression in relevant cell types and even single-cell data.

We developed a probabilistic model for personal genome interpretation, Watershed, that improves over standard methods for scoring rare variants by additionally integrating multiple transcriptomic signals from the same individual. When assessing GTEx rare variants within eQTL or sQTL colocalized regions for UKBB traits, we found that filtering by Watershed scores identified variants with larger trait effect sizes than relying only on genomic annotations. Our results demonstrate that considering predicted effects on expression could improve power for rare variant association testing by improving the quality of variant prioritization beyond what is possible using genomic annotations alone, with potential application in case/control studies. Combined, the integration of a richer set of functional rare variants into models of genetic burden and risk will improve disease gene identification and the delivery of personalized, precision genomics.

## Acknowledgements

We thank members of the Lappalainen, Mohammadi, Montgomery, and Battle labs for helpful discussions and feedback, and the artists of the graphics that we modified in Fig. 1A, found at http://www.allvectors.com/human-organs/. We also thank J. Bonnie for providing comments on the manuscript and K. Tayeb for reviewing code.

## Author’s contributions

NMF, BJS, JE, PM, SBM, and AB designed the study, performed analyses, and wrote the manuscript. NMF, XL and SBM conducted eOutlier analysis and NMF, EKT, JRD and SBM conducted tissue-specific eOutlier analysis. BJS and AB developed SPOT and conducted sOutlier analysis. JE, PM, BK and TL conducted aseOutlier analysis. BJS and AB developed Watershed. NMF, CS, and SBM conducted trait analyses. FA and KGA generated processed expression, splicing, and cis-eQTL data. SEC generated ASE call sets. ANB generated sQTL colocalizations. YP generated eQTL colocalizations. ATH and TLA performed MVP lookups. AJS and IH generated structural variant data. CS, EKT, JRD, and TL provided feedback on the manuscript.

## Data and code availability

The data analyzed for this study are available to authorized users via dbGaP under accession phs000424.v8 and on the GTEx portal (http://gtexportal.org/). ANEVA-DOT is accessible at https://doi.org/10.5281/zenodo.3406690. SPOT is available at https://github.com/BennyStrobes/SPOT. Code for correlation eOutlier calls can be found at https://github.com/nmferraro5/correlation_outliers. Watershed is available at https://github.com/BennyStrobes/Watershed.

