## Supplementary Materials for "Diverse transcriptomic signatures across human tissues identify functional rare genetic variation"

#### Materials and Methods

**GTEx data**

All human donors were deceased, and informed consent was obtained via next-of-kin consent for the collection and banking of deidentified tissue samples for scientific research. The research protocol was reviewed by Chesapeake Research Review Inc., Roswell Park Cancer Institute's Office of Research Subject Protection, and the institutional review board of the University of Pennsylvania. We used the RNA-sequencing, allele-specific expression, and whole-genome sequencing (WGS) data from the v8 release of the GTEx project and assessed expression data across the 49 biological tissues with at least 70 samples. Sample size varied across tissues, with average missingness of ~50%. Self-reported ancestry for these individuals spanned three of the major continental populations with the majority (n=714 with WGS) comprising individuals of predominantly European ancestry, 121 individuals with African ancestry, 11 with Asian ancestry, and 12 unknown or other. The generation of these data are described in the supplementary information of (The GTEx Consortium 2019, in submission).

**Rare variant annotations**

We retained all SNVs and indels that passed quality control in the GTEx VCF, variant calling described in (The GTEx Consortium 2019, in submission), using the hg38 genome build. Structural variants were called according to (*33*) on the subset of individuals available from V7 with GenomeSTRiP (*34*) GSCNQUAL set to limit the false discovery rate (FDR) for each variant type. Genome STRiP's IntensityRankSumAnnotator was used to evaluate FDR based on available Illumina Human Omni 5M gene expression array data. GSCNQUAL was limited to ≥ 1 for GenomeSTRiP deletions and ≥ 8 for multi-allelic copy number variants, corresponding to an FDR of 10%. The GSCNQUAL cutoff for GenomeSTRiP duplications was set at ≥ 17, the point where the FDR plateaued at 15.1% and did not fluctuate more than ±1% for over 50 steps in increasing GSCNQUAL score. Additionally, the Mobile Element Locator Tool (MELT) version 2.1.4 (*35*) was run using MELT-SPLIT to identify ALU, SVA, and LINE1 insertions into the test genomes. MELT calls that were categorized as "PASS" in the VCF info field, had an ASSESS score ≥ 3, and SR count ≥ 3 were retained. SV calls were then lifted to the hg38 genome build using liftOver from the Genome Browser (*36*).

We defined rare variants as those with < 1% MAF within GTEx and, for SNVs and indels, also occurring at < 1% frequency in non-Finnish Europeans within gnomAD (*37*). Novel variants were those that occurred in GTEx but were not found in gnomAD. GTEx singletons had an average allele frequency of 0.0030 in gnomAD and doubletons had an average frequency of 0.0096.

Annotation of protein-coding regions and transcription factor binding site motifs was generated by running Ensembl VEP (version 88). Loss of function (LoF) annotation was generated using loftee. Conservation scores (Gerp, PhyloP, PhastCons) were downloaded from UCSC genome browser and CADD scores were extracted from a pre-compiled annotation file (https://cadd.gs.washington.edu/download) using variant scores from the hg38 genome build.

**Expression outlier calling**

Within each tissue, we log2-transformed the expression values ($\log_2(\text{TPM} + 2)$), where TPM is the number of transcripts per million mapped reads, generated by RNA-SeQC (*38*) using the GENCODE v26 gene annotation, available through the GTEx portal. We subsetted to autosomal lincRNA and protein-coding genes and restricted to genes with at least 6 reads and TPM > 0.1 in at least 10% of individuals. We scaled the expression of each gene to mean of 0 and standard deviation of 1 to avoid the deflation of outlier values caused by quantile normalization. As we expected unmeasured technical confounders to impact expression, for each tissue we estimated hidden factors for the transformed expression matrix using PEER (*39*). The number of PEER factors retained was based on sample size and matched the values chosen in the GTEx eQTL analyses (The GTEx Consortium 2019, in submission), which were 15 for sample sizes less than or equal to 150, 30 for less than 250, 45 for less than 350, and 60 otherwise. We obtained expression residuals by regressing out PEER factors, the top three genotype principal components, sex, and the genotype of the strongest cis-eQTL per gene in each tissue using the following linear model:

$$Y_g = \mu_g + \sum_{n=1}^{N} \alpha_{g,n} P_n + \sum_{k=1}^{3} \beta_{g,k} G_k + \gamma_g S + \delta_g Q + \varepsilon_g$$

where $Y_g$ is the transformed expression of gene $g$, $\mu_g$ is the mean expression level for the gene, $P_n$ is the nth PEER factor, $G_k$ are the top k genotype principal components, $S$ is the sex covariate, and $Q$ is the genotype of the strongest cis-eQTL for gene $g$. We then re-scaled the expression residuals $\varepsilon_g$ for each gene, to obtain corrected expression Z-scores for each individual per gene per tissue.

For each gene, we calculated an individual's median Z-score across all tissues for which data were available, restricting to individuals with measurements in at least five tissues. To account for situations where widespread extreme expression might occur in an individual due to non-genetic influences, we excluded 39 individuals where the proportion of tested genes that were outliers at a threshold of |median Z-score| > 3 exceeded 1.5 times the interquartile range of the distribution of proportion outlier genes across all individuals. We then use the median Z-scores per individual to determine eOutliers and used a threshold of 3 to determine the outlier set of genes. Controls were defined as any individual with a |median Z-score| of less than 3 (or another threshold as indicated) for the same set of genes as those with any outlier individual. We allowed a gene to have multiple outlier individuals and an individual could be an outlier for multiple genes. Code for generating outlier calls was modified from the scripts available at https://github.com/joed3/GTExV6PRareVariation.

**Split read count quantification and processing**

LeafCutter (*40*) provided an annotation-free approach for RNA splicing quantification allowing us to capture split reads overlapping rare exon-exon junctions. Junctions were extracted from

WASP-corrected BAM files with a modified version of the “bam2junc.sh” script from LeafCutter that only retained reads that passed WASP filters (The GTEx Consortium 2019, in submission). Next in each tissue separately, junction reads were clustered using the “leafcutter_cluster.py” script from LeafCutter, with the options “--maxintronlen 500000” and “mincluratio 0”. LeafCutter assigns exon-exon junctions into mutually exclusive sets, termed clusters. Each exon-exon junction in a cluster had to share a splice site with at least one other exon-exon junction in that cluster, but could not share a splice site with an exon-exon junction from another cluster. A cluster had to contain at least two exon-exon junctions.

Next, in each tissue separately, we applied the following series of custom filters to the LeafCutter results in order to remove exon-exon junctions with low expression while retaining rare exon-exon junctions:

1. Removed exon-exon junctions where no sample has >= 15 split reads
2. Re-defined LeafCutter cluster assignments after removal of exon-exon junctions (according to the above filter) and removed exon-exon junctions that no longer shared a splice site with any other exon-exon junction.
3. Removed all exon-exon junctions belonging to a LeafCutter cluster where more than 10% of the samples had less than 3 reads summed across all exon-exon junctions assigned to that LeafCutter cluster.

Next, we merged LeafCutter cluster assignments across all 49 tissues to make a specific LeafCutter cluster comparable across multiple tissues. For this, we re-defined LeafCutter cluster assignments using the union of all exon-exon junctions that passed the above filters across 49 tissues. Lastly, we mapped our LeafCutter clusters to genes by intersecting splice sites, defining a Leafcutter cluster with splice sites of annotated exons. We limited to genes used in expression outlier calling (described in “Expression outlier calling” section). If an annotated splice site was in a LeafCutter cluster, we considered the LeafCutter cluster mapped to the gene. It was therefore possible for a LeafCutter cluster to map to multiple genes. We filtered LeafCutter clusters, and their corresponding exon-exon junctions, to those that were mapped to at least one gene. Finally, we removed any LeafCutter clusters with more than 20 exon-exon junctions due to computational limitations of SPOT.

**SPOT: Overview**

sOutliers were identified separately for each LeafCutter cluster in each tissue using Splicing Outlier deTection (SPOT). For a given LeafCutter cluster in a given tissue, we defined a matrix, $X$ (dim NxJ), where each row corresponds to one of $N$ samples, each column corresponding to one of $J$ exon-exon junctions, and each element was the number of raw split read counts corresponding to that row’s sample and that column’s exon-exon junction. We were able to compute a p-value representing how abnormal a given sample’s splicing patterns were for the given LeafCutter cluster as follows:

1. Fitted parameters of Dirichlet-Multinomial distribution based on observed data $X$ in order to capture the distribution of split read counts mapping to this LeafCutter cluster

2. Used fitted Dichlet-Multinomial distribution to compute the Mahalanobis distance for each of the N samples
3. Computed Mahalanobis distance for 1,000,000 samples simulated from the fitted Dirichlet-Multinomial and use these 1,000,000 Mahalanobis distances as an empirical distribution to assess the significance of the $N$ real Mahalanobis distances

**SPOT: Dirichlet-Multinomial parameter estimation**
We defined a Dirichlet-Multinomial (DM) probability distribution based on data from $N$ samples to capture the probability that a split read would map to each of the $J$ junctions in the Leafcutter cluster:
Let $x_{nj}$ be the raw number of split reads mapped to the j[th] junction in the n[th] sample and $t_n = \sum_{j=1}^{J} x_{nj}$ be the total number of split reads mapped to any junction in this LeafCutter cluster in the nth sample. Then
$x_{n1}, ..., x_{nJ} \mid t_n \sim DM(t_n, \alpha_1 p_1, \ldots, \alpha_J p_J)$ where $p_j = \frac{exp(c_j)}{\sum_{k=1}^{J} exp(c_k)}$
We used the following non-informative Gamma prior distribution to stabilize optimization:
$\alpha_j \sim Gamma(1 + 1e^{-4}, 1e^{-4})$
We then performed maximum likelihood estimation (via LBFGS as implemented in STAN) to learn the optimal parameter settings of $\alpha_1, \ldots, \alpha_J$ and $c_1, \ldots, c_J$ ( $\hat{\alpha}_1, \ldots, \hat{\alpha}_J$ and $\hat{c}_1, \ldots, \hat{c}_J$ ) from the N samples. We were able to also deterministically compute the optimal values of each $p_j$ ( $\hat{p_j}$ ) from each $\hat{c}_j$ .

**SPOT: Mahalanobis distance**
The Mahalanobis distance is the multivariate generalization of how many standard deviations a point is from the mean taking into account the covariance structure. After learning the parameters of the Dirichlet-Multinomial distribution for a specific LeafCutter cluster (ie $\hat{\alpha}_1, \ldots, \hat{\alpha}_J$ and $\hat{c}_1, \ldots, \hat{c}_J$ ; see “SPOT calling: Dirichlet-Multinomial parameter estimation”), we were able to compute the mean vector ( $\mu_n$ ) and covariance matrix ( $\Sigma_n$ ) for a specific sample $n$ , according to the Dirichlet-Multinomial. Using $\mu_n$ and $\Sigma_n$ we were able to compute the Mahalanobis distance of sample $n$ $(MD_n)$ . The covariance matrix of the Dirichlet-Multinomial ( $\Sigma_n$ ) is of rank $J - 1$ because one of the dimensions is always a linear combination of the other $J - 1$ dimensions. As such, we approximated $\Sigma_n^{-1}$ with the pseudo-inverse of $\Sigma_n$ when computing the Mahalanobis distance.

**SPOT: Empirical distribution to assess significance**
For a given LeafCutter cluster, we have already computed the Mahalanobis distance of each of the $N$ samples according to the fitted Dirichlet-Multinomial distribution for that LeafCutter cluster. However, the Mahalanobis distance is biased by the dimensionality of the space (i.e. the number of junctions assigned to the LeafCutter cluster). In order to convert the Mahalanobis distance to a test statistic that was not biased by dimensionality, we simulated an empirical

distribution of Mahalanobis distances for each LeafCutter cluster. Specifically, for one LeafCutter cluster we drew 1,000,000 random samples from the fitted Dirichlet-Multinomial distribution assuming each of these random samples has 20,000 reads mapped to the LeafCutter cluster $(t_n = 20000)$. We then computed the Mahalanobis distance of each of these 1,000,000 samples and used the 1,000,000 Mahalanobis distances as an empirical distribution that converted our N Mahalanobis distances (from the real data) into p-values.

**SPOT: Gene level correction**

To compute a splicing outlier p-value for a gene associated with $C$ LeafCutter clusters, we first computed minimum p-value across all $C$ clusters for the gene. However, the minimum of a list of p-values is not a valid p-value. To address this, we computed the probability of observing a minimum p-value according to a probability density function defining the minimum across $C$ independent uniform random variables between 0 and 1:

$$p(min(pvalue_1, \; ... \; , \; pvalue_C) \; <= \; z) \; = \; 1 \; - \; (1-z)^C$$

This approach made the conservative, simplifying assumption that all clusters mapped to a gene were independent of one another.

We excluded individuals (global outliers) where the proportion of tested genes that were outliers (at a threshold of median p-value < .0027) exceeded 1.5 times the interquartile range of the distribution of proportion outlier genes across all individuals.

**ASE outlier calling**

Allelic expression (ASE) data was produced as described in (Castel, et al, 2019, in submission). We used the Analysis of Expression VAriation Dosage Outlier Test (ANEVA-DOT; (*16*)) to identify genes in each individual that showed an excessive imbalance of ASE, relative to the population. Briefly, ANEVA-DOT relies on tissue-specific estimates of genetic variation in gene dosage, $V^G$, derived by Analysis of Expression VAriation (ANEVA) on a reference population ASE data to identify genes in individual test samples that are likely affected by rare variants with unusually large regulatory effects. We calculated reference $V^G$ estimates from GTEx v8 data from 15,201 RNA-seq samples spanning 49 tissues and 838 individuals with WGS data (16, 17). Across all analyzed tissues we estimated $V^G$ a total of 2,727,867 times using all available autosomal aeSNPs (variants used to assess allelic expression) with at least 30 reads in 6 individuals. These estimates are publicly available at https://github.com/PejLab/Datasets/tree/master/Reference_Vg_Estimates. We used the ANEVA-DOT tool R package (https://doi.org/10.5281/zenodo.3406690) to calculate a p-value for every gene-individual pair with allelic expression data and a corresponding $V^G$ estimate (Fig S5). The p-value can be interpreted as the result of a binomial test of allelic imbalance, that is overdispersed for each gene individually according to its expected dosage variation in a given tissue in the population. Genes with significant ANEVA-DOT p-values are referred to as aseOutliers in this text. We tested all tissues available for each GTEx v8 individual, using only genes with a minimum coverage of 8 reads spanning an aeSNP and with $V^G$ estimates available (49 tissues, median genes per tissue = 4899, Fig S4). For each gene expressed we considered the aeSNP with the highest coverage in an individual.

For all single-tissue analyses, we removed global outlier genes and individuals from each tissue group independently, based on the lists of ASE blacklisted genes and individuals available at https://github.com/PejLab/Datasets/tree/master/ANEVA_DOT_frequencies. These genes are likely to have poor $V^G$ estimates due to the presence of different ASE patterns within the gene or other global biological factors. Global outlier lists were derived from lists of FDR-corrected p-values. In all analyses unless otherwise specified, we did not apply an FDR control procedure and instead imposed a higher threshold for declaring significance, to be consistent with expression and splicing outliers. For cross-tissue analyses, we calculated median ANEVA-DOT p-values for genes which were expressed in more than 5 tissues, without removing known global outliers first. Therefore, to account for genes with poor $V^G$, we applied the filtering steps described in (*16*) on the resulting individual-level median p-values. Briefly, we removed individuals with too few genes tested (*n*<Q1-1.5IQR), removed individuals with too many outliers (*n*>Q3+1.5IQR), and removed genes which appeared as outliers too many times across individuals with a score available (genes that are likely to be called as outliers in more than 1% of cases, Fig S4) . For consistency with the other outlier detection methods we declared significance at $p < 0.0027$, equivalent to $|Z| > 3$.

**Correlation-aware expression outlier calling**

We subsetted to a set of individuals and tissues with < 75% missingness, leading to 762 individuals and 29 tissues. We imputed missing expression values to improve our estimate of the tissue-by-tissue covariance matrix per gene that would be used in outlier calling. We used K-nearest neighbors in the impute R package (*41*) with k = 200 to impute values for missing tissues per individual on a gene by gene basis. We chose the value of k by comparing reconstruction error across k = [1, 5, 10, 15, 20, 25, 30, 35, 40, 45, 50, 55, 60, 65, 70, 80, 90, 100, 200, 300] on a set of 1000 randomly selected genes with 5% of individuals held-out for evaluation. We tested several other potential imputation methods and saw similar performance (Fig S16), which included a multivariate normal expectation-maximization (EM), mean imputation (MEAN), soft thresholded iterated SVD imputation (SVD), and penalized matrix decomposition (PMD). For these additional imputation methods, we used the following parameters, determined in the same way as described above: EM - max iterations = 3 and tolerance = $1 \times 10^{-6}$, SVD - lambda = 20 and rank = 20, and PMD - lambda = 1 and rank = 5.

From the imputed matrix, we estimated the tissue covariance matrix, $\widehat{\Sigma}$, for each gene. We calculated the Mahalanobis distance for each individual-gene pair as follows:

$$d^2 = x_g^T \widehat{\Sigma}_g^{-1} x_g,\ d^2 \sim \chi_p^2$$

Where $x_g$ is the vector of observed expression values for gene $g$ across tissues, $\widehat{\Sigma}_g$ is the estimated covariance matrix for gene $g$. We assigned a p-value to each gene-individual from the chi-squared distribution with degrees of freedom $p$ equal to the number of tissues available for that individual. We used a two step correction procedure, first correcting via Bonferroni for all genes tested within an individual and then applying Benjamini-Hochberg correction across all tests with p < 0.0027. When assessing nearby rare variant enrichments, we removed genes that had an extreme number of outlier individuals, based on 3*IQR, as compared to the total set of tested genes. For the set of tissue-specific correlation outliers, we subsetted to outliers driven by a single tissue, requiring remaining available tissues for that individual to have a |Z-score| < 2 for the outlier gene.

**Enrichment calculations**

We calculated relative risk enrichments as the proportion of outliers with a given variant type nearby the outlier gene over the proportion of non-outlier individuals with the given variant type nearby the same set of genes. We included 95% confidence intervals estimated via a normal approximation. When assessing rare variant enrichments overall and by category, we used a 10kb +/- window around the gene body. When considering variant categories per outlier, if more than one rare variant was present nearby the outlier gene, we assigned each gene-individual to a single variant category based on the following ordering: duplications (DUP), copy number variations (CNV), deletions (DEL), breakend (BND), inversions (INV), transposable elements (TE), splice, frameshift, stop, transcription start site (TSS), conserved non-coding, coding, or other non-coding, and subsetted to the 527 individuals with structural variant calls. Unless otherwise specified, we used a threshold of median p-value < 0.0027 (chosen to match |median Z-score| > 3) to define outliers. When considering variants in different windows upstream from the gene, we constructed exclusive distance ranges from each gene, beginning with the gene body + 10kb window used previously, and then we intersected rare variants with windows 1bp-200kb, 200kb-400kb, 400kb-600kb, 600kb-800kb, and 800kb-1Mb upstream from the set of outlier genes.

**Alternative splicing enrichment calculations**

We performed several enrichment analyses specific to sOutliers. For all of these analyses, we used sOutlier calls at the LeafCutter cluster level (instead of the gene level) in order to get more accurate enrichments. We excluded individuals identified as global outliers at the gene level (see "SPOT: gene level correction"). We also used a stringent median p-value threshold of $1 \times 10^{-5}$ to identify outlier clusters and limited enrichment analysis to SNVs.

1. **Relative risk of rare variant in window around splice site**. We computed the relative risk of rare variants being located at various windows around splice sites for outlier clusters relative to non-outlier clusters (splice enrichment 1a). For example, if the window was [0,2], we mapped a variant to a cluster if that variant were less than or equal to two base pairs away from observed donor and acceptor splice sites ([D-2, D+2] and [A-2, A+2] based on notation in Fig 2C) for that cluster. Relative risk was then calculated as the proportion of outlier (LeafCutter cluster, individual) pairs with a mapped rare variant over the proportion non-outlier (LeafCutter cluster, individual) pairs with a mapped rare variant, while limiting analysis to LeafCutter clusters with at least one

outlier individual. We included 95% confidence intervals estimated via a normal approximation.

2. **Relative risk of rare variant at position relative to splice site.** We first mapped rare variants to clusters if the rare variants were less than or equal to 1000 base pairs from an observed donor or acceptor splice site ([A-1000, A+1000] and [D-1000, D+1000] based on notation in Fig 2C). We then mapped each variant to its nearest splice site in that cluster and calculated its position relative to that splice site. Then, to compute the positional relative risk at position D-1 (for example), we computed the fraction of outlier variants mapped to a donor splice site that were at position D-1 over the fraction of non-outlier variants mapped to a donor splice site that were at position D-1. We added a constant of 1 to all counts in the contingency table to stabilize enrichments. We included 95% confidence intervals estimated via a normal approximation.
3. **Junction Usage for splicing median p-value outliers**. We used the "junction usage" statistic to quantify whether an individual used a splice site more or less than the background population. A positive junction usage value intuitively means the individual uses the splice site more than the background population, while a negative junction value means an individual uses a splice site less than the background population. More concretely to compute the junction usage for an individual $i$ and junction j, we first computed the following ratio in each tissue (in which that individual $i$ is expressed) separately: $\frac{Fraction\ of\ reads\ in\ cluster\ mapping\ to\ junction\ j\ for\ individual\ i}{Fraction\ of\ reads\ in\ cluster\ mapping\ to\ junction\ j\ for\ non-outliers\ individuals}$
We added a constant of 1 to the above contingency table to stabilize enrichments. The "junction usage" statistic is simply the natural logarithm of the median of the above statistic across all tissues in which individual $i$ is expressed.

**Enrichment of outlier pairs within a given window**

To test if nearby genes were more likely to share outlier status, we counted how many times two consecutive genes within a given genomic distance (defined based on the gene start position) in a given individual were both reported outliers. We considered multi-tissue outliers and analyzed each class of outliers independently. To derive the expected number of such occurrences, for each individual we used sampling without substitution to produce a random set of genes of the same size. Samples were drawn from a list of all genes that had been reported as an outlier at least once across all methods to avoid skewing the statistic by genes never reported as outliers. The expected value for each given window size was derived by averaging over all individuals. To ensure the stability of enrichment estimates at each window size, the sampling process was repeated until Monte Carlo error dropped below 10% of the expected number of outlier co-occurrences. For sOutliers this procedure was repeated once with all outlier genes included and once after removing 80 genes sharing a cluster with another outlier gene, see "Split read count quantification and processing".

We annotated all outliers occurring in a given window with the set of nearby rare variants for each gene in the pair. For each included variant category, defined above, we calculated a relative risk by taking the proportion of outlier pairs within the window for which one or both genes had a rare variant in that category near the gene over the proportion of control individuals for which the same was true for the same gene, restricting to individuals with genetic data available. We included 95% confidence intervals estimated via a normal approximation, and we defined controls as individuals who were outliers for only one of the genes in the outlier pair.

**Single-tissue rare variant enrichment**

We tested for enrichment of rare variants near single-tissue gene expression outliers using the same variant list and relative risk enrichment definition as for cross-tissue outliers and with all individuals with both an expression outlier score and genotype information available. Under this definition of an eOutlier, a gene is only considered in one tissue at a time, i.e. without aggregating the gene's score across all tissues in an individual where it is expressed. Among ASE and splicing outliers, we removed tissue-specific global outlier genes prior to performing enrichment analysis. We converted expression Z-scores to a two-tailed *z*-test p-value for direct comparison to the other outlier methods. We tested for enrichment of rare variants at increasingly stringent significance thresholds for each individual tissue, then reported the range of enrichment scores across all tissues, separated by outlier type and significance threshold.

**Tissue-specific enhancer enrichments**

We obtained tissue-specific enhancer annotations for 12 tissues from Epigenomics Roadmap (*42*) and mapped to GTEx tissues (Table S1). We subsetted to the tissue-specific correlation outliers that occurred in one of the 12 mapped tissues. To calculate the relative risk of a rare variant, including both SNVs and indels, in a tissue-matched enhancer, we took controls as all individual-gene pairs that were not correlation outliers and randomly assigned them to the same set of tissues as in the outlier group, matched by gene. We used any enhancer region annotated to a given tissue within a 500kb window around the outlier gene to capture the majority of potential enhancers, which can act at longer distances (*18*). We calculated matched enhancer enrichments as the proportion of tissue-specific outliers for which a rare variant fell within a nearby tissue-matched enhancer over the proportion of control individuals for which the same was true. For unmatched enhancer enrichments, we calculated the proportion of tissue-specific outliers with a rare variant falling in any tissue-specific enhancer region across the 12 tissues considered, without regard to the tissue driving the outlier call, within a 500kb window over the proportion of controls with a rare variant in any enhancer region within the same window of the same gene set.

**Watershed model overview**

Watershed is a hierarchical Bayesian model that predicts regulatory effects of rare variants on a specific outlier signal based on the integration of multiple transcriptomic signals along with genomic annotations describing the rare variants. Watershed models instances of (gene, individual) pairs to predict the regulatory effects of rare variants nearby the gene. The Watershed model for a particular (gene, individual) pair, assuming $K$ outlier signals, consists of three layers (Fig 4a):

1. A set of variables **G** $= G_1, ..., G_P$ representing the P observed genomic annotations aggregated over all rare variants in the individual that are nearby the gene.

2. A set of binary latent variables **Z** = $Z_1, ..., Z_K$ representing the unobserved functional regulatory status of the rare variants on each of the K outlier signals. Let $Z^S$ be the set of all possible values that **Z** can take on. The size of $Z^S$ is $2^K$.
3. A set of categorical nodes **E** = $E_1, ..., E_K$ that represents the observed outlier status of the gene for each of the K outlier signals. We allow for missingness in **E**.

A fully connected conditional random field (CRF) (*43*) is defined over variables $Z$ given $G$, where we let W represent the set edges among $Z$. Variables $E_i$ are each connected only to the corresponding latent variable $Z_i$. Specifically, the following conditional distributions together define the full Watershed model:

A. $Z \mid G \sim CRF(\alpha, \beta_1, ..., \beta_k, \theta)$
B. $E_k \mid Z_k \sim$ Categorical $(\phi_k)$ $\forall\ k \in K$
C. $\phi_k \sim$ Dirichlet $(C, ..., C)$
D. $\beta_k \sim$ Normal $(0, \frac{1}{\lambda})$

where,

- $\beta_k \in R^P\ \forall\ k \in K$ are the parameters defining the contribution of the genomic annotations to the CRF for each outlier signal ($k$)
- $\alpha \in R^K$ are the parameters defining the intercept of the CRF for each outlier signal ($k$)
- $\theta \in R^{(K choose 2)}$ are the parameters defining the edge weights between pairs of outlier signals (Notational note: $\theta_{tq} = \theta_{qt}$)
- $\phi_k \forall\ k \in K$ are the parameters defining the categorical distributions of each outlier signal
- $C$ and $\lambda$ are hyper-parameters of the model

Explicitly, our CRF probability distribution is defined as:

$$P(Z \mid G, \beta_1, ..., \beta_K, \alpha, \theta) = exp(\sum_{k \in K} \alpha_k Z_k + \sum_{(t,q) \in W} \theta_{tq} Z_t Z_q + \sum_{k \in K} \beta_k G\, Z_k - A(G, \theta, \beta_1, ..., \beta_K)$$

where $A(G, \theta, \beta_1, ..., \beta_K) = log(\sum_{Z^* \in Z^S} exp\,(\sum_{k \in K} \alpha_k {Z^*}_k + \sum_{(t,q) \in W} \theta_{tq} {Z^*}_t {Z^*}_q + \sum_{k \in K} \beta_k G\, {Z^*}_k))$

Because **Z** is unobserved, the Watershed log-likelihood objective over instances $n = 1, ..., N$:

$$\sum_{n=1}^{N} log \sum_{Z^* \in Z^S} P(E_n, G_n, Z^* \mid \beta_1, ..., \beta_K, \alpha, \theta, \phi_1, ..., \phi_K)$$

is non-convex. We therefore optimize model parameters using Expectation-Maximization (EM) as described in the following sections.

**Watershed exact inference optimization routine**

When the number of outlier signals ( $K$ ) is small (an approximate rule being 4 or less), Watershed parameters can be optimized using exact inference updates within EM as follows:

In the E-step for instances $n = 1, ..., N$ : we compute posterior distributions over the latent variables ( $Z^{(n)}$ ), conditioned on the current model parameters ( $\beta_1, ..., \beta_K, \alpha, \theta, \phi_1, ..., \phi_K$ ) and the observed data ( $G^{(n)}$ and $E^{(n)}$ ). For example, the joint posterior probability of $Z^{(n)} = Z$ for the nth instance can be computed as:

$$\omega^{(n)}(Z^{(n)} = Z) = exp(\sum_{k \in K}(\alpha_k Z_k + \beta_k G^{(n)} Z_k + I(E_k^{(n)}) log(P(E_k^{(n)} | Z_k)) + \sum_{(t,q) \in W} \theta_{tq} Z_t Z_q$$
$$- A(G^{(n)}, E^{(n)}, \theta, \beta, \alpha, \theta, \phi)$$

$$A(G^{(n)}, E^{(n)}, \theta, \beta, \alpha, \phi) = log(\sum_{Z^* \in Z^S} exp(\sum_{k \in K}(\alpha_k Z^*_k + \beta_k G^{(n)} Z^*_k + I(E_k^{(n)}) log(P(E_k^{(n)} | Z^*_k))$$
$$+ \sum_{(t,q) \in W} \theta_{tq} Z^*_t Z^*_q))$$

where,
$I(E_k^{(n)})$ is an indicator function for whether $E_k^{(n)}$ is observed. Given the joint posterior probability distribution, we can marginalize (sum over) specific dimensions (outlier signals) to obtain:

1. Marginal posterior distributions for each dimension $i$ (where $Z^W$ is the set of all possible values that **Z** can take on excluding dimension $i$ ):

$$\omega^{(n)}_{single}(Z_i) = \sum_{Z^* \in Z^W} \omega^{(n)}(Z^*)$$

2. Pairwise marginal posterior distributions for each pair of dimensions $i,j$ (where $Z^W$ is the set of all possible values that **Z** can take on (excluding dimension $i$ and dimension $j$ )):

$$\omega^{(n)}_{pair}(Z_i, Z_j) = \sum_{Z^* \in Z^W} \omega^{(n)}(Z^*)$$

Both the marginal posterior distributions and the pairwise marginal posterior distributions are used in the M-step as follows. We update $\beta$ , $\alpha$ , and $\theta$ by optimizing the conditional random field as follows:

$$argmax_{\beta, \alpha, \theta} \sum_{n=1}^{N} \sum_{Z^* \in Z^S} log(P(Z^* | G^{(n)}, \beta, \alpha, \theta)) \, \omega^{(n)}(Z^*) - \tfrac{\lambda}{2}||\beta||_2 - \tfrac{\lambda}{2}||\theta||_2$$

Here $\lambda$ is an L2 penalty hyper-parameter derived from the Gaussian priors on $\beta$ and $\theta$ . We optimized this objective function by running L-BFGS on the closed-form gradient updates.

In the second part of the M-step, we update $\phi_k \forall k \in K$ as follows:

$$\phi_k(s, t) = \sum_{n=1}^{N} I(E_k^{(n)} = t) \, \omega^{(n)}_{single}(Z_k^{(n)} = s) + C$$

where,
$I$ is an indicator operator, $t$ is the categorical value of expression $E_k^{(n)}$, $s$ is the possible binary values of $Z_k^{(n)}$, and $C$ is the hyperparameter based on the Dirichlet prior on $\phi$.

Once the EM algorithm has converged, we use the marginal posterior distributions for each dimension $i$ in each instance $n$ ($\omega^{(n)}_{single}(Z_i = 1)$) as estimates of probability that the nth (gene, individual) pair has a nearby variant that has a functional effect on the gene (with respect to outlier dimension i).

**Watershed approximate inference optimization routine**
When the number of outlier signals ($K$) is large (an approximate rule being 5 or more), it becomes computationally intractable to optimize Watershed parameters using exact inference updates, so we use approximate inference updates within EM as follows:

For the E-step, we wish to compute approximate estimates of the following posterior probability distribution:

$$\omega^{(n)}(Z^{(n)} = Z) = exp(\sum_{k \in K}(\alpha_k Z_k + \beta_k G^{(n)} Z_k + I(E_k^{(n)}) log(P(E_k^{(n)} | Z_k)) + \sum_{(t,q) \in W} \theta_{tq} Z_t Z_q - A(G^{(n)}, E^{(n)}, \theta, \beta, \alpha, \theta, \phi)$$

$$A(G^{(n)}, E^{(n)}, \theta, \beta, \alpha, \phi) = log(\sum_{Z^* \in Z^S} exp(\sum_{k \in K}(\alpha_k Z^*_k + \beta_k G^{(n)} Z^*_k + I(E_k^{(n)}) log(P(E_k^{(n)} | Z^*_k)) + \sum_{(t,q) \in W} \theta_{tq} Z^*_t Z^*_q))$$

To approximate this function $\omega^{(n)}(Z^{(n)})$, we use the Mean-Field Approximation (a subclass of Variational Inference) (*44*) and optimize $q^{(n)}(Z^{(n)})$ to minimize the KL-divergence between $q^{(n)}(Z^{(n)})$ and $\omega^{(n)}(Z^{(n)})$
where,

$$q^{(n)}(Z^{(n)}) = \prod_{k \in K} q_k^{(n)}(Z_k^{(n)}) \text{ where } q_k^{(n)}(Z_k^{(n)}) = (\mu_k^{(n)})^{z_k^{(n)}}(1 - \mu_k^{(n)})^{(1 - z_k^{(n)})}$$

To minimize the KL-divergence for a given sample $n$, we perform coordinate descent on each $\mu_k^{(n)}$ while holding all other dimensions (values of $\mu_j^{(n)}$) constant. Given that $N(k)$ represents the set of all nodes that share an edge with node $k$, the variational update for each $\mu_k^{(n)}$ is then:

$$\mu_k^{(n)(update)} = \frac{exp(a_k + I(E_k^{(n)}) log(P(E_k^{(n)} | Z_k{=}1)))}{exp(I(E_k^{(n)}) log(P(E_k^{(n)} | Z_k{=}0)) + exp(a_k + I(E_k^{(n)}) log(P(E_k^{(n)} | Z_k{=}1)))} \text{ where } a_k = \alpha_k + \beta_k G^{(n)} + \sum_{j \in N(k)} \theta_{kj} \mu_j^{(n)}$$

More specifically, for one instance $n$, we iteratively do the following until convergence:

1. Loop through all $K$ dimensions in a random order, and update each $\mu_k^{(n)}$ given the most recent values of $\mu_j^{(n)} \forall j \in N(k)$. Since coordinate ascent is not guaranteed to reach the global optimum, we used damped updates for each $\mu_k^{(n)} \forall k \in K$ in order to decrease the chance of getting stuck at a local optimum:
    a. $\mu_k^{(n)(iter\ i+1)} = (1 - \eta) * \mu_k^{(n)(iter\ i)} + (\eta) * \mu_k^{(n)(update)}$

b. We use a damping value ( $\eta$ ) of 0.8.

2. Compute the average difference, across all $K$ dimensions, between the values of $\mu_k^{(n)}$ from the current iteration and values of $\mu_k^{(n)}$ from the previous iteration. Converge if the average difference is less than 1x $10^{-8}$.

Using the same notation as in "Watershed exact inference optimization routine", Mean Field allows us to approximate the following expectations using converged estimates of $\mu_k^{(n)}$:

1. $\omega^{(n)}(Z^{(n)}) \approx \prod_{k\in K} (\mu_k^{(n)})^{z_k^{(n)}}(1-\mu_k^{(n)})^{(1-z_k^{(n)})}$
2. $\omega^{(n)}_{pair}(Z_i^{(n)}, Z_j^{(n)}) \approx (\mu_i^{(n)})^{z_i^{(n)}}(1-\mu_i^{(n)})^{(1-z_i^{(n)})}\,(\mu_j^{(n)})^{z_j^{(n)}}(1-\mu_j^{(n)})^{(1-z_j^{(n)})}$
3. $\omega^{(n)}_{single}(Z_i^{(n)}) \approx (\mu_i^{(n)})^{z_i^{(n)}}(1-\mu_i^{(n)})^{(1-z_i^{(n)})}$

We use both the approximate marginal posterior distributions and the approximate pairwise marginal posterior distributions in the M-step. However, when the number of dimensions ( $K$ ) is large, optimization of the parameters ( $\beta$ , $\alpha$ , and $\theta$ ) defining the conditional random field becomes intractable. Therefore, we approximated the CRF objective function with the Pseudolikelihood (*45*) of the CRF. Given variational estimates of $\mu_i^{(n)}(Z_i^{(n)})$ for all values of dimensions ( $i$ ) and all samples ( $n$ ), the (log) Pseudolikelihood objective function (including priors on coefficients) is given by:

$$\sum_{n=1}^{N}\sum_{k\in K}\left(\alpha_k\mu_k^{(n)} + \beta_k G^{(n)}\mu_k^{(n)} + \sum_{j\in N(k)}\theta_{kj}\mu_k^{(n)}\mu_j^{(n)} - A(k, n, \theta, \beta, \alpha)\right) - \tfrac{\lambda}{2}\|\beta\|_2 - \tfrac{\lambda}{2}\|\theta\|_2$$

$$A(k, n, \theta, \beta, \alpha) = log\left(\sum_{z=0}^{1} exp\left(\alpha_k z + \beta_k G^{(n)} z + \sum_{j\in N(k)}\theta_{kj}\, z\, \mu_j^{(n)}\right)\right)$$

We computed closed form gradient updates of the above objective function and then optimized it using L-BFGS.

In the second part of the M-step, we update $\phi_k \forall\ k \in K$ as follows:

$$\phi_k(s, t) = \sum_{n=1}^{N} I(E_k^{(n)} = t)\, \omega^{(n)}_{single}(Z_k^{(n)} = s) + C$$

Where $I$ is an indicator operator, $t$ is the categorical value of expression $E_k^{(n)}$, $s$ is the possible binary values of $Z_k^{(n)}$, and $C$ is the hyperparameter based on the Dirichlet prior on $\phi$.

Once the EM algorithm has converged, we use marginal posterior distributions for each dimension i, in each instance n ( $\omega^{(n)}_{single}(Z_i = 1)$ ) as estimates of probability that the nth (gene, individual) pair has a nearby variant that has a functional effect on the gene (with respect to outlier dimension $i$ ).

**GAM and RIVER**

The genomic annotation model (GAM) is L2-regularized logistic regression using genomic annotations (**G**) as features and the binary outlier status of a specific outlier signal as the response variable. One GAM model was trained for each outlier signal.

The only difference between Watershed and RIVER is that in RIVER $\theta$ is fixed to be a vector of zeros. This allows RIVER to be optimized precisely as described in "Watershed exact inference optimization routine" assuming $\theta$ is fixed to be zero. It is important to note that RIVER has changed slightly since its initial development (*15*) in the following way: we now use a categorical distribution ( $\phi$ ) with three categories instead of two to model $E \mid Z$ . This change in RIVER was made in order to make it directly comparable to Watershed.

**Applying Watershed to jointly model ASE, splicing, and expression**

We first applied Watershed to the GTEx V8 data using 3 outlier signals: median ASE, splicing, and expression. Recall, Watershed requires a set of genomic annotations (**G**) and a corresponding set of categorical outlier signals (**E**) over (gene, individual) instances. We first limited to a set of (gene, individual) pairs that passed the following set of filters in all 3 outlier signals:

1. The individual was not a global outlier
2. The gene has measured outlier signal for the corresponding individual
3. The gene has at least one individual that is an outlier (median p-value < .01)

This yielded a set of 36,702 (gene, individual) pairs that we used for training and evaluating the Watershed framework.

To generate the genomic annotations (**G**) for each (gene, individual) pair, we limited to SNVs that fell within the gene body or +/- 10kb of each of the gene and then extracted 47 genomic annotations (Table S4) describing each of the SNVs including regulatory element annotations, conservation scores, and derived genomic scores from other models such as CADD. If a (gene, individual) pair had more than one SNV mapped to the gene, the genomic annotations were aggregated across the SNVs with simple transformations to generate gene-level genomic annotations (Table S4). The resulting gene-level genomic annotations were standardized (mean 0 and standard deviation 1) before running Watershed. $1.93 \times 10^{-5}$

We generated the categorical outlier signals (**E**) for each (gene, individual) pair using 3 categories per outlier signal. It is important to note that because of the filters described above there is no missingness in **E**. For aseOutliers and sOutliers, we assigned a gene with median p-value ( $p$ ) to:

1. Category 1 if $-log_{10}(p + 10^{-6}) < 1$
2. Category 2 if $1 <= -log_{10}(p + 10^{-6}) < 4$
3. Category 3 if $-log_{10}(p + 10^{-6}) >= 4$

For eOutliers, we assigned a gene with median p-value ( $p$ ) and median Z-score (z) to:

1. Category 1 if $-log_{10}(p + 10^{-6}) > 1$ and z < 0

2. Category 2 if $-log_{10}(p + 10^{-6}) <= 1$
3. Category 3 if $-log_{10}(p + 10^{-6}) > 1$ and z > 0

To train and evaluate Watershed, we identified the 3,411 cases where two or more individuals had the same rare SNV(s) near a particular gene. We held out those instances and trained Watershed on the remaining instances. For training, we set the hyperparameter $C$ equal to 30, motivated by the number of training instances. To select the hyperparameter $\lambda$, we trained and evaluated GAM on the training data for each outlier signal independently(assigning a sample an outlier label if outlier p-value < .01) with 5-fold cross validation while running a gridsearch on $\lambda$ =.1,.01,.001. We selected the $\lambda$ with the largest median area under the precision recall curve (AUPRC) across the 5 folds. Each precision recall curve aggregated predictions across the three outlier signals. The optimal $\lambda$ was found to be 0.001. Before running Watershed, we initialized $\alpha_k$ and $\beta_k$ to be the intercept and slope parameters, respectively, of GAM (when $\lambda = 0.001$) trained on the full training data for outlier signal $k$. $\theta$ was initialized to all zeros. $\phi_k$ was initialized using the MAP updates described in “Watershed exact inference optimization routine”, except we used the GAM (when $\lambda = 0.001$) posterior probabilities to approximate $\omega^{(n)}_{single}(Z_k^{(n)} = s)$.

We evaluated various trained models (Watershed, RIVER, GAM, CADD) using the 3,411 cases where two individuals had the same rare SNV(s) near a particular gene (we will refer to these instances as N2 pairs). Specifically, we estimated the posterior probability of a functional rare variant (according to each of the models) in the first individual from the pair, allowing Watershed to use all data available for that individual. We then used the outlier status of the second individual as a ‘label’ for evaluation. In order to make the fraction of outlier instances comparable between different outlier signals, we defined a (gene, individual) pair to be an outlier for a specific outlier signal if its outlier p-value was ranked amongst the 1% most significant p-values for that outlier signal (across training and N2 pair instances). For an N2 pair, we did this evaluation in both directions: predict on the first individual and evaluate on the second, as well as predict on the second individual and evaluate on the first. Importantly, none of the N2 pairs were used in training any of the models.

**Applying Watershed to jointly model outlier signals from each tissue (tissue-specific Watershed)**

Next, we trained three independent tissue-specific Watershed models (one each for ASE, splicing, and expression) where each model considered effects in all tissues, giving 49 phenotypes, corresponding to 49 Z and E variables each. In order for these models to be comparable to the model described in “Applying Watershed to jointly model three outlier types”, we used the same set of (gene, individual) pairs. We therefore used the same extracted and processed genomic annotations (**G**).

We generated the categorical outlier signals (**E**) for each (gene, individual) pair in a particular tissue (for a particular outlier signal) using 3 categories. It is important to note that, unlike the first application of Watershed to three median signals, there is now missingness in **E** as a (gene, individual) pair does not have measured outlier signal across all 49 tissues in GTEx. For ASE and splicing outliers, for a particular tissue, we assigned a gene with p-value ($p$) to:

1. Category 1 if $-log_{10}(p + 10^{-6}) < 1$
2. Category 2 if $1 <= -log_{10}(p + 10^{-6}) < 4$
3. Category 3 if $-log_{10}(p + 10^{-6}) >= 4$

For expression, outliers, for a particular tissue, we assigned a gene with p-value ($p$) and Z-score (z) to:

1. Category 1 if $-log_{10}(p + 10^{-6}) > 1$ and z < 0
2. Category 2 if $-log_{10}(p + 10^{-6}) <= 1$
3. Category 3 if $-log_{10}(p + 10^{-6}) > 1$ and z > 0

To train and evaluate Watershed, we identified the 3,411 cases where two individuals had the same rare SNV(s) near a particular gene. We held out those instances and trained Watershed on the remaining instances. For training, we set the hyperparameter $C$ equal to 10, motivated by the number of training instances with observed outlier calls. We selected $\lambda = 0.001$ based on cross-validation in "applying Watershed to jointly model three outlier types". We initialized $\alpha_t$ and $\beta_t$ to be the intercept and slope parameters, respectively, of GAM (when $\lambda = 0.001$) trained on the full training data from tissue *t*. $\theta$ was initialized to all zeros. $\phi_t$ was initialized using the MAP updates described in "Watershed exact inference optimization routine", except we used the GAM (when $\lambda = 0.001$) posteriors to approximate $\omega^{(n)}{}_{single}(Z_k^{(n)} = s)$.

We took a very similar approach as described in "Applying Watershed to jointly model ASE, splicing and expression" to evaluate various trained models (Watershed, RIVER, GAM). In this setting however, both model predictions and outlier labels were in a single tissue as opposed to the median across tissues. As **E** contains missingness in this setting, we required both individuals in the N2 pair to have observed outlier signal for the gene of interest in the corresponding tissue.

##### UKBB GWAS

We assessed GWAS summary statistics from the UK Biobank (UKBB) phase 2, as made available by the Neale lab (http://www.nealelab.is/uk-biobank/). We subsetted the variants, either genotyped or imputed, in UKBB phase 2 to those SNVs that also appeared in any GTEx individuals and had a frequency of < 1% in GTEx, which resulted in 45,415 SNVs, filtered to those not flagged as low confidence due to very low allele counts. Because we are targeting rare variants occurring at frequencies too low to obtain a trait association with genome-wide significance, we focused on the effect size estimates and did not filter by p-value. We defined outlier variants in this context as any rare variant appearing near an eOutlier, sOutlier, or aseOutlier in GTEx and also appearing in UKBB. We defined non-outlier variants as rare GTEx variants appearing in UKBB, but not falling near an outlier of any type, though within 10kb of a gene for which any individual was an outlier. We subsetted to 34 traits tested for colocalization between the UKBB GWAS and GTEx eQTL/sQTL studies. When filtering to colocalized regions, we included as a colocalization event any gene that had a colocalization posterior probability > 0.5, for both eQTLs and sQTLs (The GTEx Consortium 2019, in submission). We combine both enloc (*46*) and coloc (*47*) results for eQTL colocalization and enloc results for sQTL colocalization. This resulted in 4,787 variants across 1,323 genes and 34 traits with any significant co-localization in an included UKBB trait (Table S2). We transformed the |effect sizes| to percentiles, based on all rare GTEx SNVs that also appear in any UKBB samples tested for the included traits. When showing actual beta values for binary traits, we scaled according to the case-control ratio $\mu$ for the given trait, dividing the effect size estimates by $\mu * (1 - \mu)$.

We filtered the set of GTEx rare variants in UKBB to those in colocalized regions, defined as being in a colocalized gene or within 10kb, and by the maximum Watershed posterior for that variant-gene combination across all data types (ASE, splicing, expression) and all tested individuals. We compared this to a genomic annotation based metric, CADD. We obtain an effect size $\beta$ for both Watershed posterior and CADD score in predicting variant effect size percentiles in co-localized regions using the following model: $P \sim \beta X + \varepsilon$ , where $P$ is a vector of variant effect size percentiles and $X$ is a vector of either Watershed posteriors or CADD scores for the same variant set.

We calculated the proportion of resulting variants that fall in the top 25% of effect sizes within colocalized regions for the associated trait across a range of posterior thresholds. We compared that proportion to the set we would obtain if filtering by a CADD score chosen to return an equal number of variants, prior to intersecting with colocalized regions. Additionally, we took 1000 random samples from the set of rare variants of an equal number to the actual number obtained by filtering at each threshold and assessed the proportion of random variants that fall in the top 25% of effect sizes for each colocalized trait. For replication in the Million Veterans Program, we obtained summary statistics for a 250kb region on either side of the variant of interest for four lipid associated traits. We calculated the |effect size| percentile for all rare variants (gnomAD AF < 0.1%) in that region and plot the absolute effect sizes vs the gnomAD allele frequency.

**Watershed predictions for UKBB variants**

We used the Watershed model that was previously trained on the 34,837 (gene, individual) pairs described in “Applying Watershed to jointly model ASE, splicing, and expression” to make Watershed predictions on the 45,415 SNVs described in “UKBB GWAS”. To make genomic annotations comparable, the genomic annotations describing the 45,415 UKBB SNVs were standardized according to the mean and standard deviation of the genomic annotations from “Applying Watershed to jointly model ASE, splicing, and expression”. It is important to note that the Watershed model was trained across (gene, individual) pairs and predictions were made across (gene, SNV, individual) triplets.

### Supplementary Materials:

#### Supplementary Figures

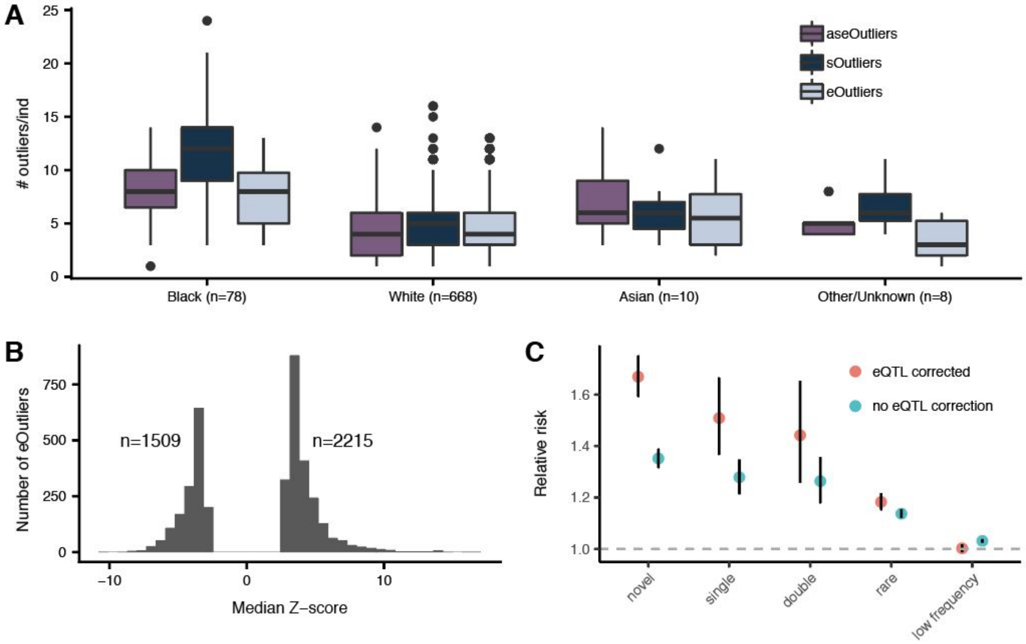


**Figure S1. Outlier distribution and effect of eQTL correction. (A)** Number of outliers per individual across each population defined by self-reported race, at a threshold of $p < 0.0027$. **(B)** Number of eOutliers split by direction of the expression effect. **(C)** Effect of correcting for strongest cis eQTL per gene on nearby rare variant (SNV+indels) enrichments for eOutliers.

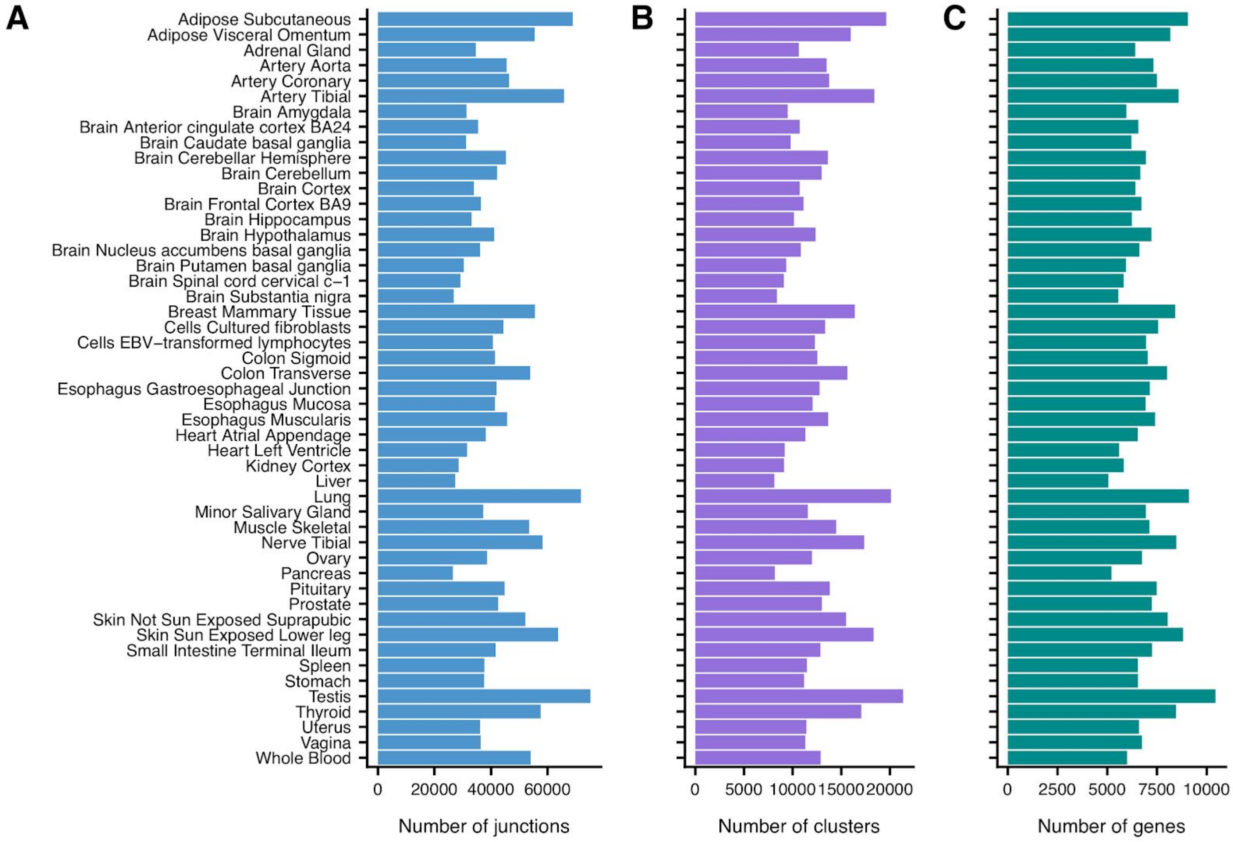


**Figure S2. sOutlier split read count processing.** The number of unique **(A)** junctions, **(B)** LeafCutter clusters, and **(C)** genes that are found in each tissue (rows) after split read count quantification and processing.

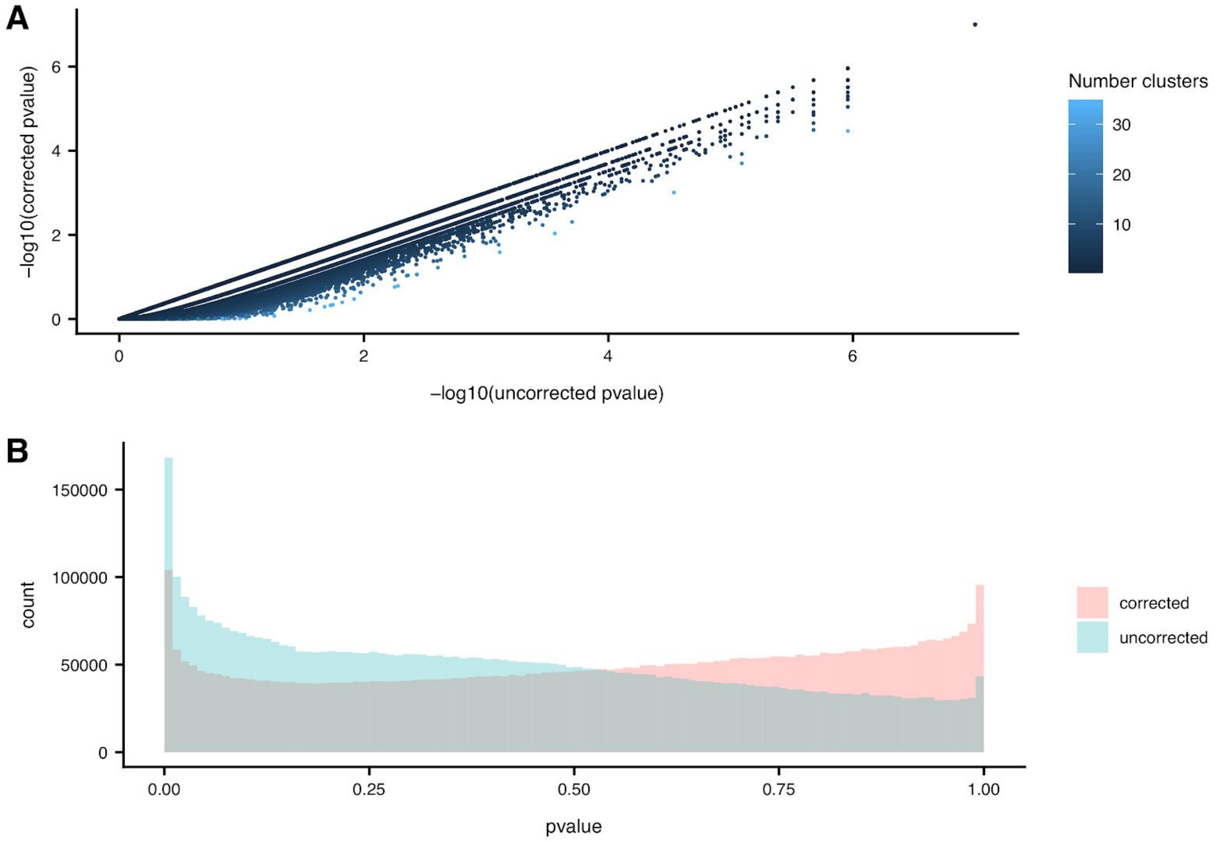


**Figure S3. SPOT gene level correction. (A)** Scatterplot showing the $-log_{10}$ sOutlier p-values in Muscle-Skeletal tissue at the gene level before the gene-level correction (x-axis) and after the gene level correction (y-axis) for the number of LeafCutter clusters mapped to each gene (color). **(B)** The distribution of sOutlier p-values in Muscle-Skeletal tissue at the gene level before the gene level correction (teal) and after the gene level correction (salmon) for the number of LeafCutter clusters mapped to each gene.

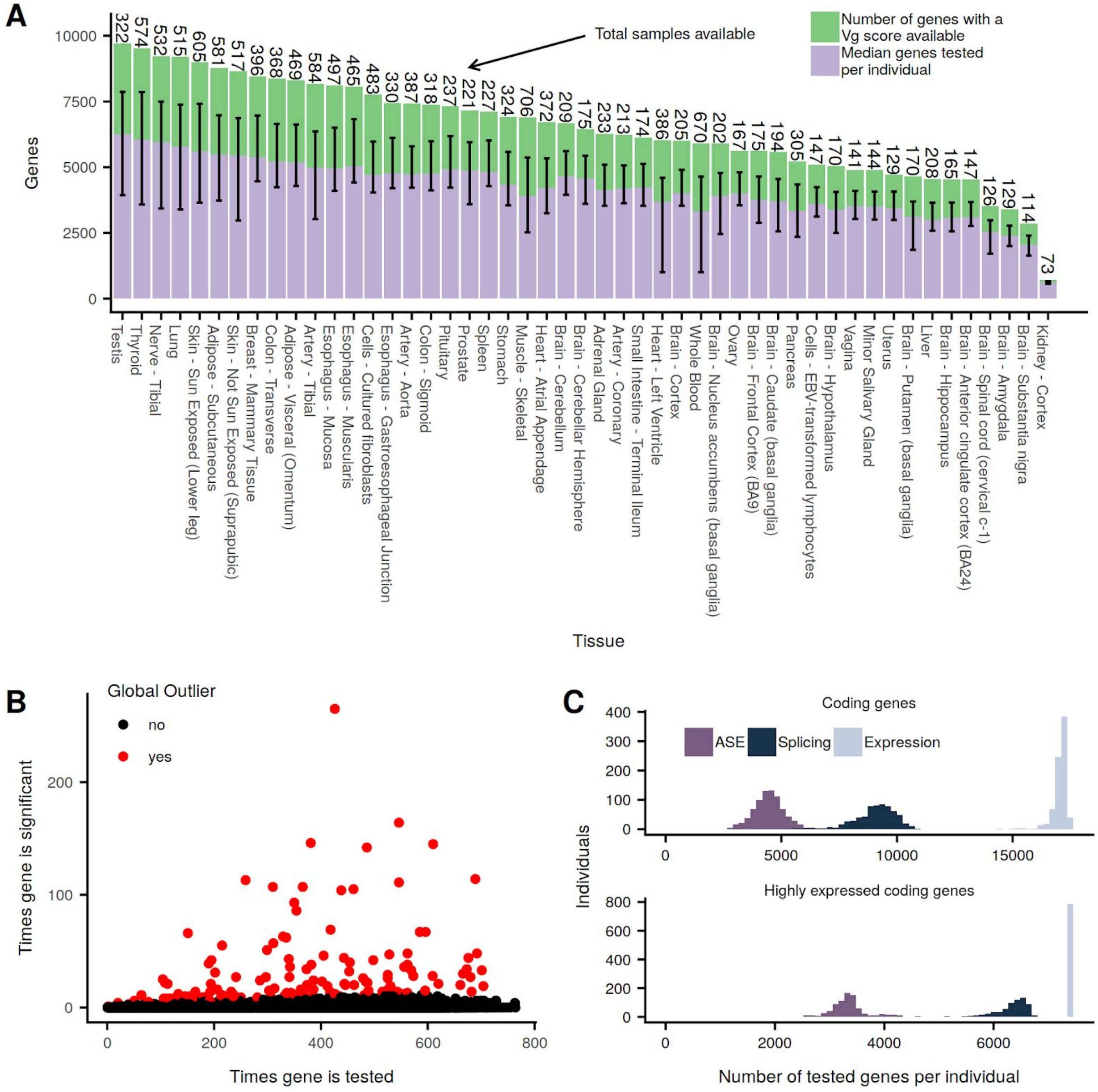


**Figure S4. Quality control for ASE processing. (A)** Average number of tests per individual tissue sample ± range. The total number of $V^G$ scores available per tissue is shown above in green, with the total samples available per tissue. **(B)** The total number of times a gene was tested by considering its median ANEVA-DOT p-value vs the number of times it was called as an outlier. We call global outliers by drawing a 95% binomial confidence interval around the outlier frequency for each gene, and flagging all genes where the interval contains 1% or greater. Global outlier genes were removed from downstream analysis. **(C)** Distribution of median number of scores available across all three outlier methods, limiting to coding genes above, and coding genes with a median TPM > 10 across all individuals and tissues below.

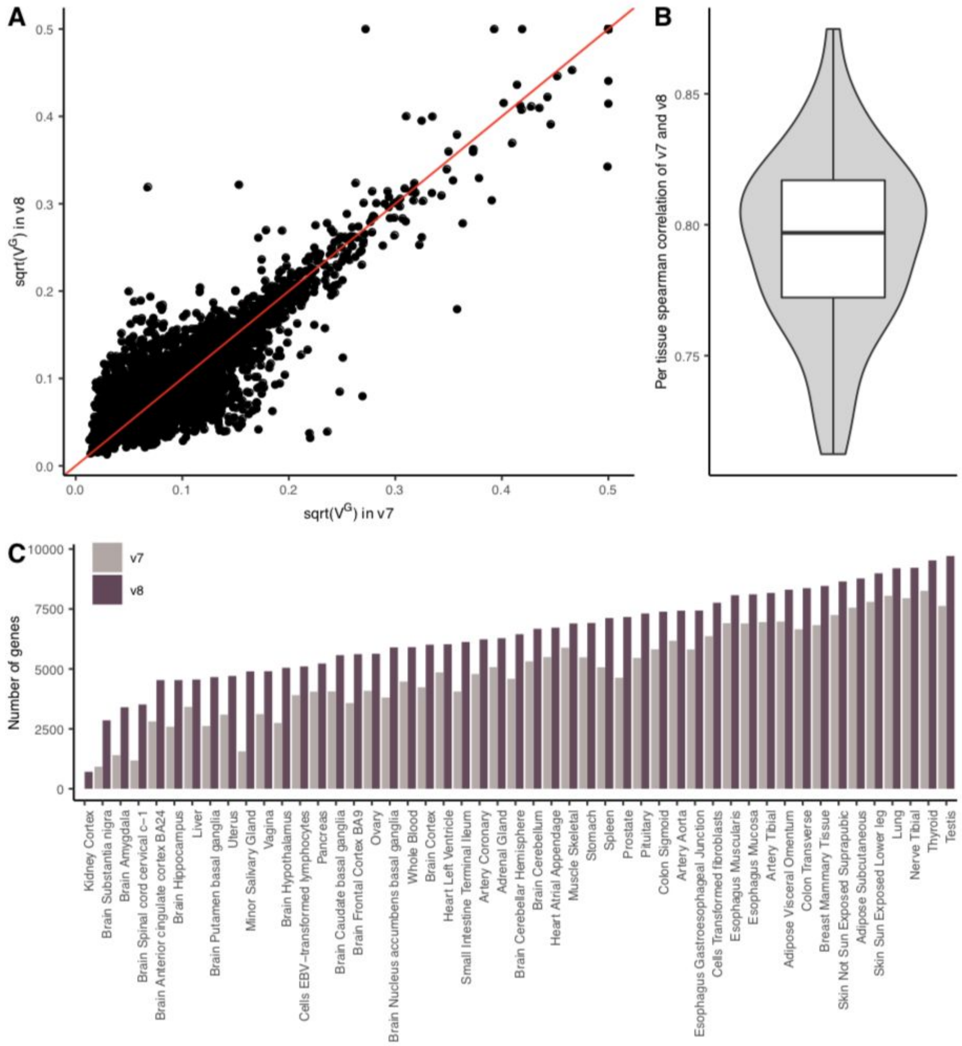


**Figure S5. ANEVA estimates of genetic variance in gene expression ($V^G$). (A)** Comparison of $V^G$ estimates for an example tissue (Adipose subcutaneous) derived from GTEx v8 dataset compared to that of v7. The red line represents x=y. **(B)** Distribution of the spearman correlation coefficient between $V^G$ estimates from v7 and v8 across all GTEx tissues. The lower and the upper whiskers indicate 1.5 interquartile range from the first and the third quartile, respectively. **(C)** The number of genes with $V^G$ estimates available across GTEx tissues in each version.

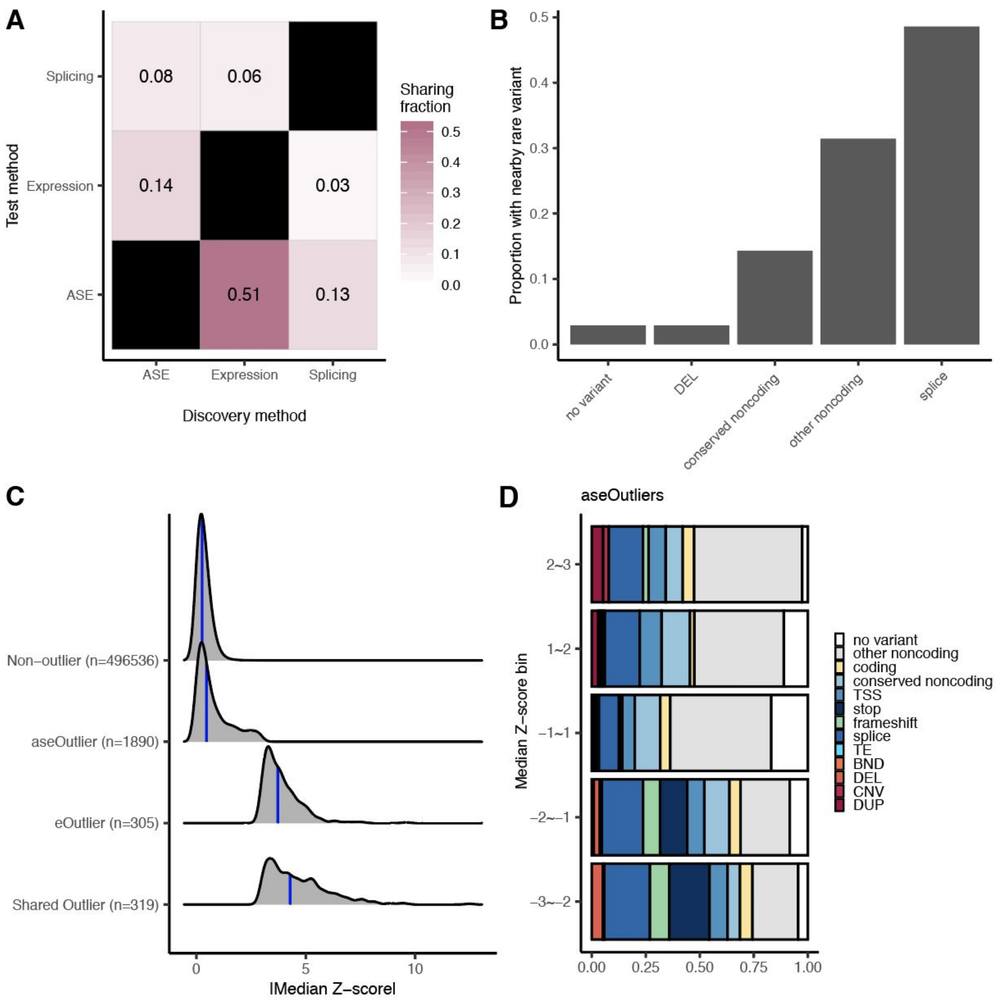


**Figure S6. Comparing outliers across methods. (A)** Of the set of individuals and genes tested across all data types, the fraction discovered via one method that also meet the outlier thresholds ($p < 0.0027$) in another method. Across all data types, 624 individuals and 8,722 genes, including 2,281,262 unique combinations, were tested by all methods. **(B)** The proportion of outliers shared across all methods assigned to the given rare variant category nearby the outlier gene. Of the 2,209 aseOutliers, 1,385 sOutliers, and 624 eOutliers discovered at this threshold among the shared set, 35 individual-gene pairs are found by all three methods, encompassing 31 unique genes. **(C)** Of the set of eOutliers and aseOutliers within this set, the distribution of |median Z-scores| for outliers in both types, expression alone, ASE alone, or non-outliers for the same set of genes. Blue lines represent the 50th percentile. **(D)** The proportion of aseOutliers with a nearby rare variant of a given type split by the corresponding median Z-score bin for the same individual-gene pair.

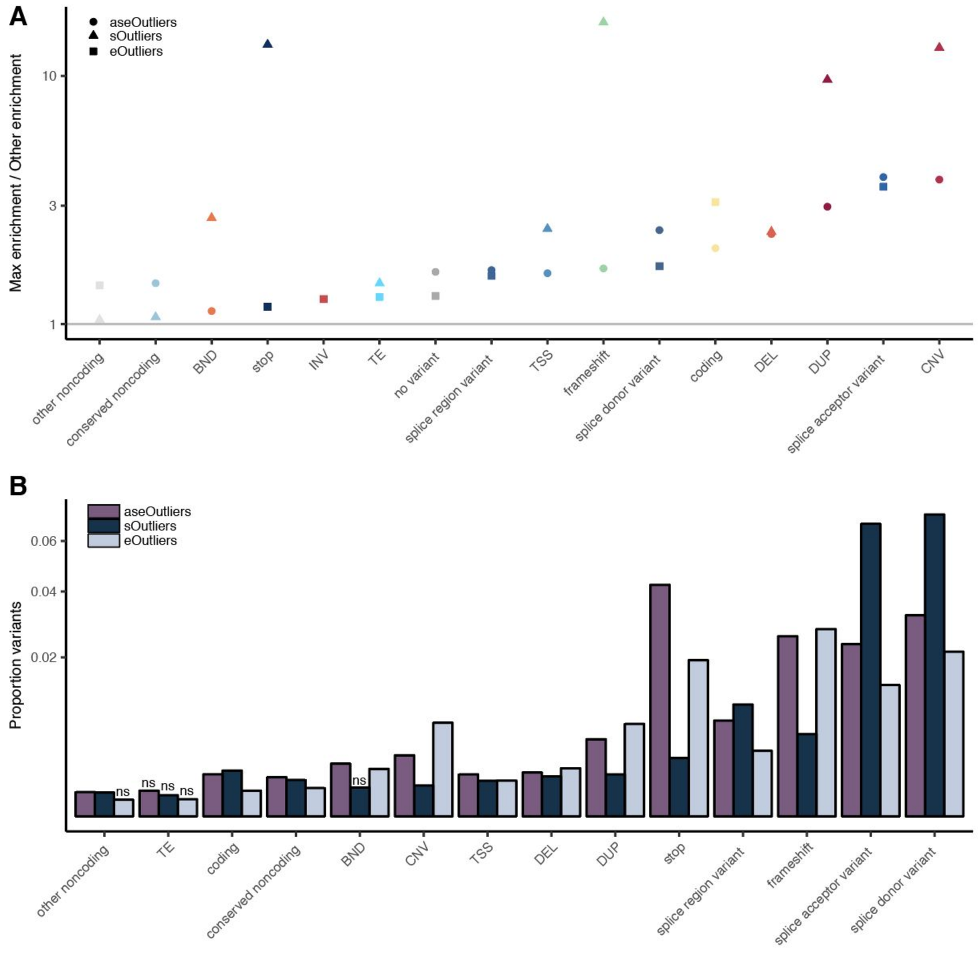


**Figure S7. Comparison of variant class enrichments across methods. (A)** For each variant category, the maximum enrichment across data types over the enrichment for the remaining two data types. **(B)** For each variant category, the proportion of variant occurrences leading to an outlier across all categories, with INV removed due to either very low or zero instances. Those marked ns indicate that in 1000 iterations permuting outlier status, a proportion greater than or equal to the actual proportion was found greater than 5% of the time.

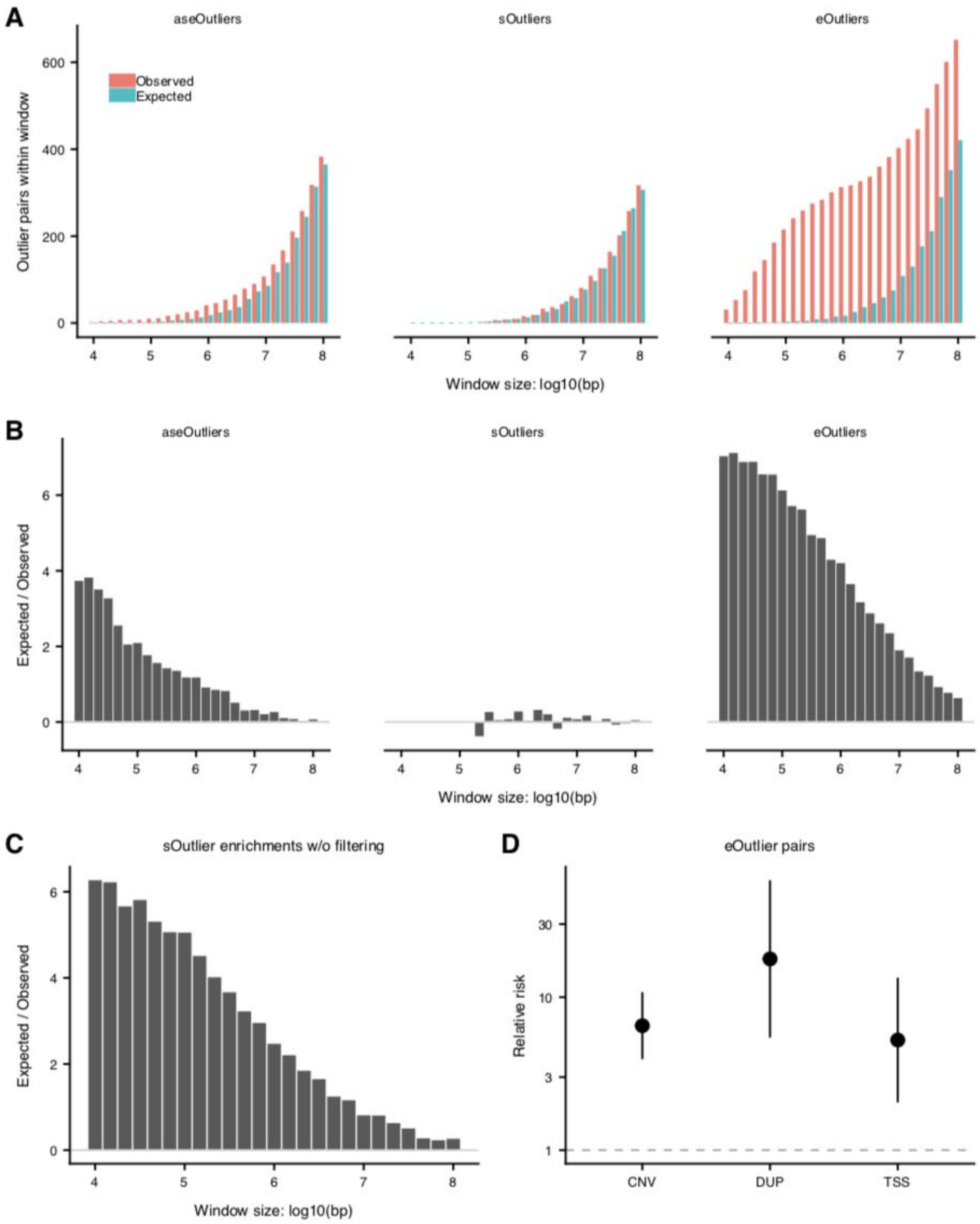


**Figure S8. Outliers occurring together within a given window. (A)** At varying window sizes, the number of observed vs expected outliers occurring together within that window. Expected numbers were generated from sampling an equal number of outlier genes from randomly chosen individuals. **(B)** The enrichment, calculated as log2 ratio of the observed number of outliers occurring in the same window over expected, across different window sizes. **(C)** In A and B, we filter out any splicing gene pairs that share a cluster, see Supplemental Methods. Here, we calculate the enrichments for sOutliers including those gene pairs. **(D)** For eOutlier pairs, the relative risk of one or both genes in the pairs found within a 100kb window having a nearby rare CNV, DUP, or TSS variant as compared to individuals who are only outliers for one of the genes in the pair.

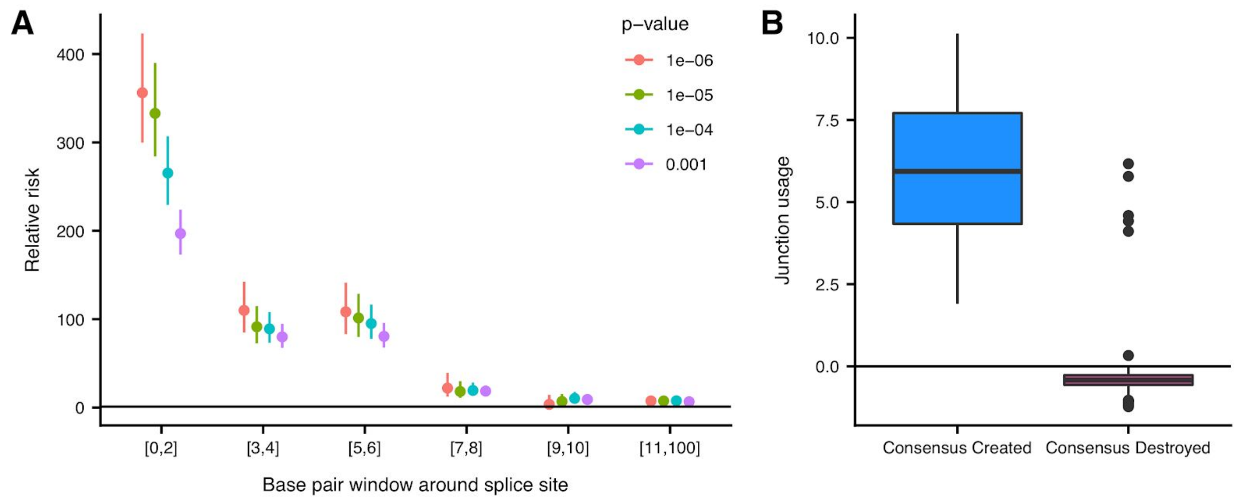


**Figure S9. Enrichment of rare variants nearby splice sites in sOutliers. (A)** Relative risk (y-axis) of rare variants within various window sizes around splice sites (x-axis) for sOutlier (median LeafCutter cluster p-value < 1 x $10^{-5}$) clusters relative to non-outlier clusters at several p-value thresholds (color). **(B)** Junction usage of a splice site is the natural log of the fraction of reads in a LeafCutter cluster mapping to the splice site of interest in sOutlier (median LeafCutter cluster p-value < 1 x $10^{-5}$) samples relative to the fraction in non-outliers samples aggregated across tissues by taking the median. Junction usage (y-axis) of the closest splice sites to rare variants that lie within the splicing consensus sequence binned by the type of variant (x-axis).

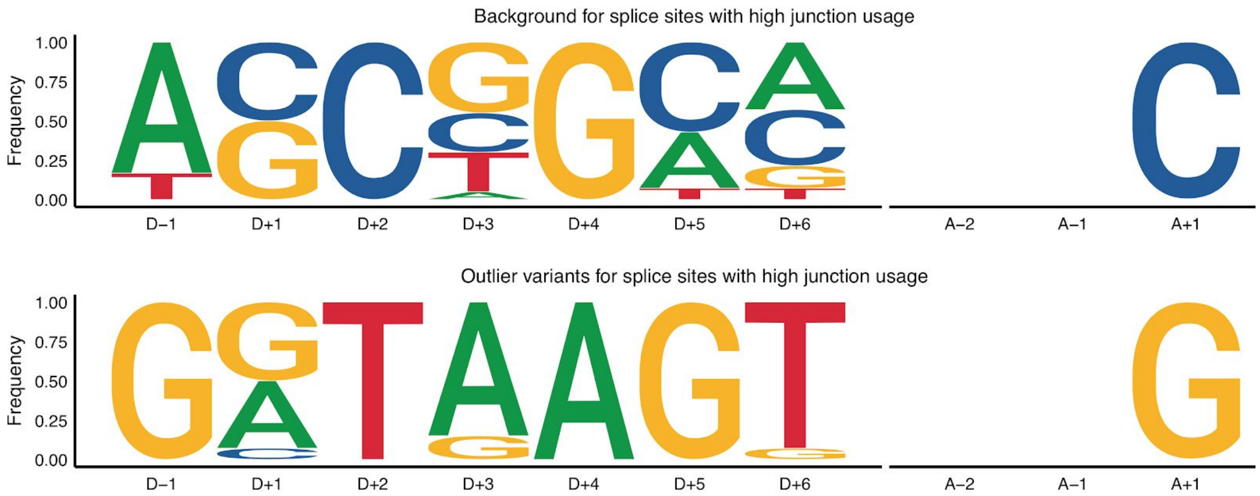


**Figure S10. sOutlier variants in consensus sequence of splice sites with high junction usage.** Independent position weight matrices showing mutation spectrums of sOutlier (median LeafCutter cluster p-value < 1 x $10^{-5}$) rare variants at positions relative to splice sites with positive junction usage (ie. splice sites used more in outlier individuals than in non-outliers).

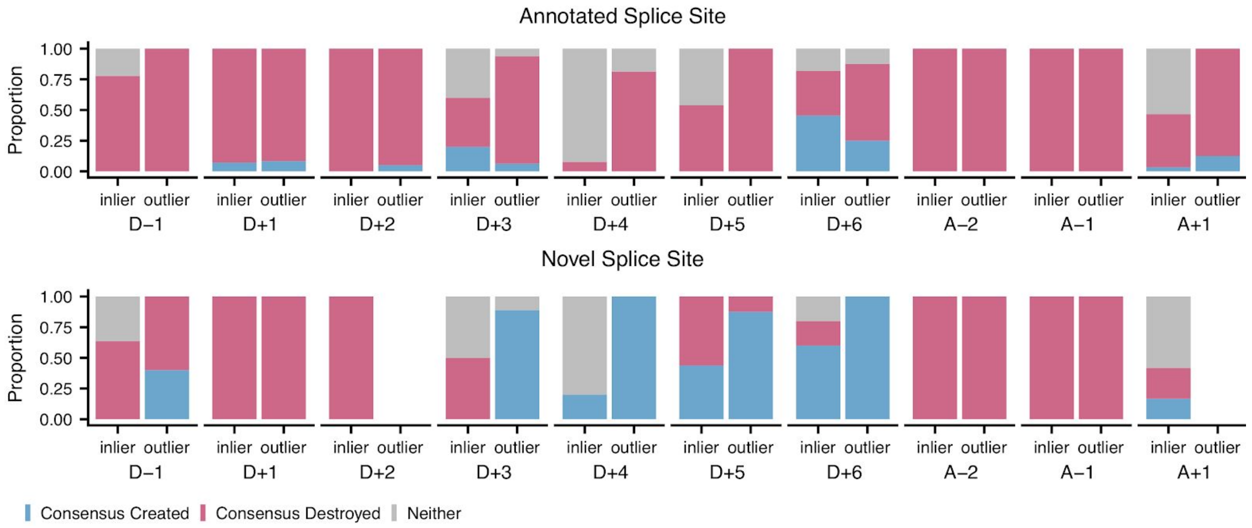


**Figure S11. sOutlier variants in consensus sequence of annotated and novel splice sites.** Proportion of sOutlier (median LeafCutter cluster p-value < 1 x $10^{-5}$) and non-outlier variants, at each position in the splicing consensus sequence, that create the consensus sequence (blue) or destroy the consensus sequence (red) where variants are binned by whether the nearby splice site is annotated or novel (rows).

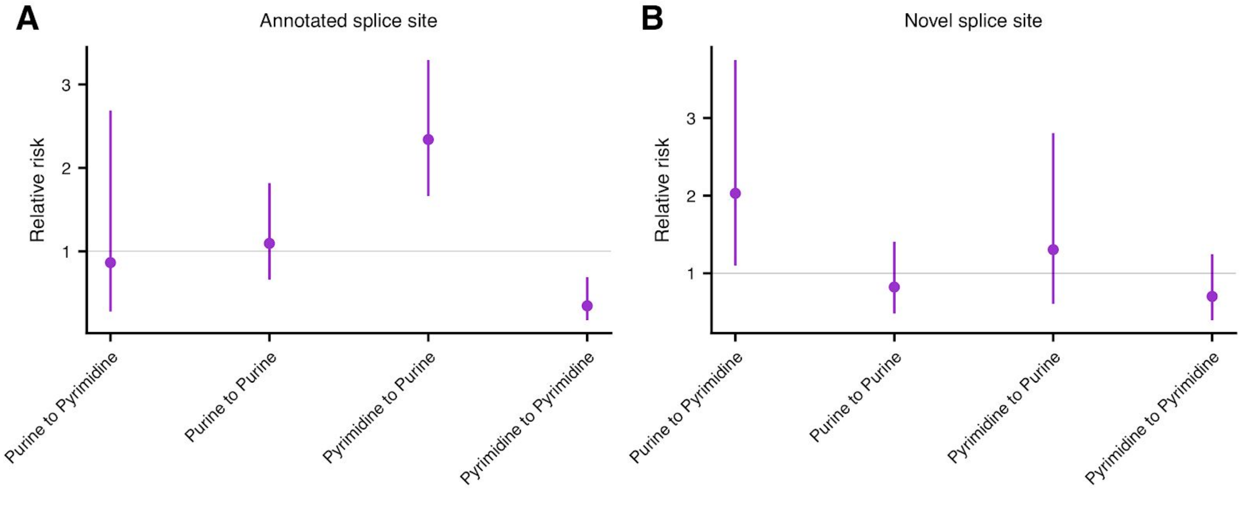


**Figure S12. sOutlier variant type enrichments in PPT.** Relative risk for sOutliers relative to non-outliers (median LeafCutter cluster p-value < 1 x $10^{-5}$) of having a rare variant that is located in PPT (5 to 35 base pairs upstream from an acceptor splice site) having a specific mutation spectrum (x-axis). Relative risk calculation done separately for annotated **(A)** or novel **(B)** splice sites.

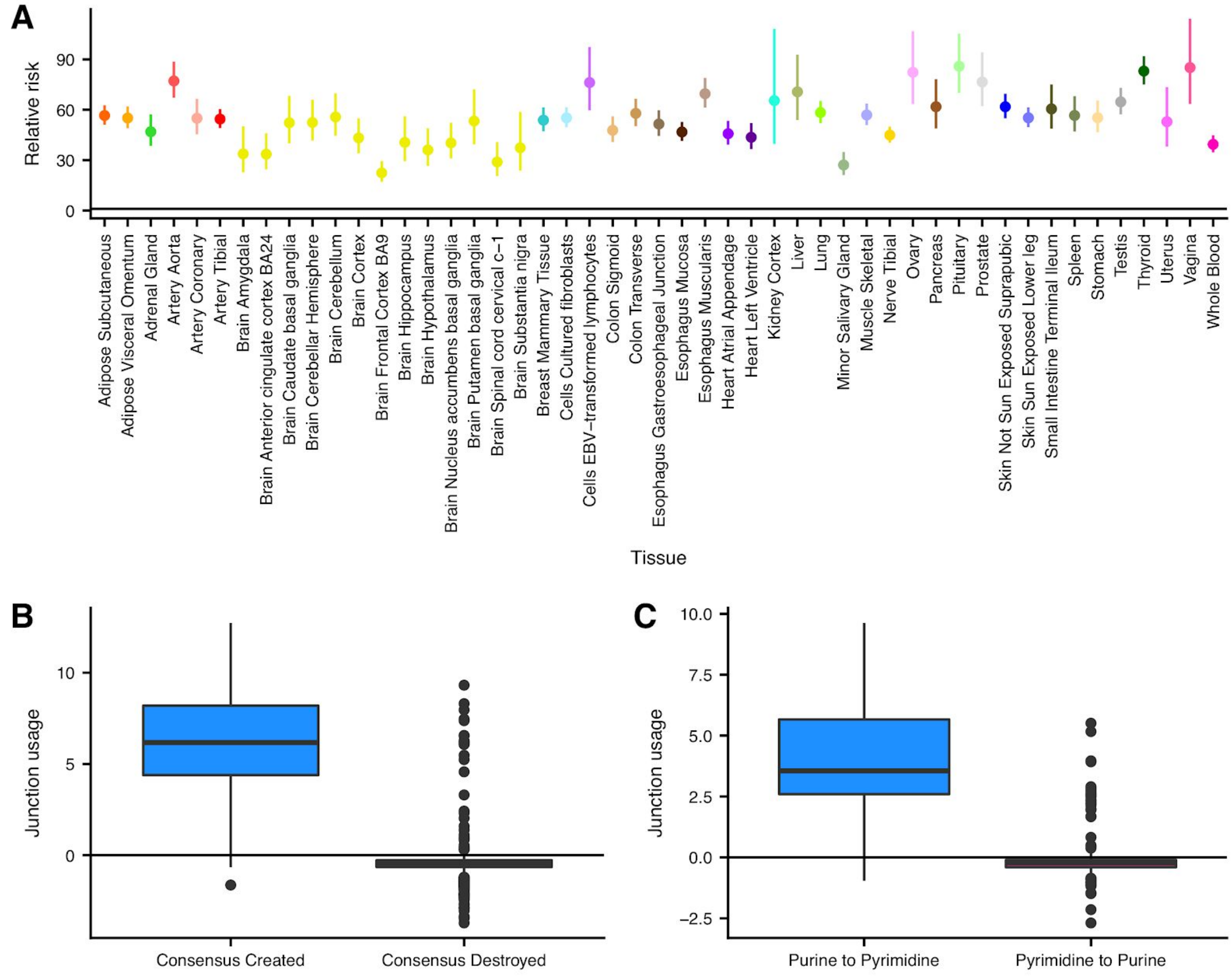


**Figure S13. Single tissue sOutlier enrichments. (A)** Relative risk, in each tissue independently, of rare variants being located in a 6 base pair window around splice sites for sOutlier LeafCuter clusters (per tissue LeafCutter cluster p-value < 1 x $10^{-5}$) relative to non-outlier clusters. **(B, C)** Per tissue junction usage of a splice site is the natural log of the fraction of reads in a LeafCutter cluster mapping to the splice site of interest in sOutlier (per tissue LeafCutter cluster p-value < 1 x $10^{-5}$) samples relative to the fraction in non-outliers samples, in a single tissue. **(B)** Per tissue junction usage (y-axis) of the closest splice sites to rare variants that lie within the splicing consensus sequence binned by the type of variant (x-axis). **(C)** Per tissue junction usage (y-axis) of the closest splice sites to rare variants that lie within a PPT ([A-5, A-35]) binned by the type of variant (x-axis).

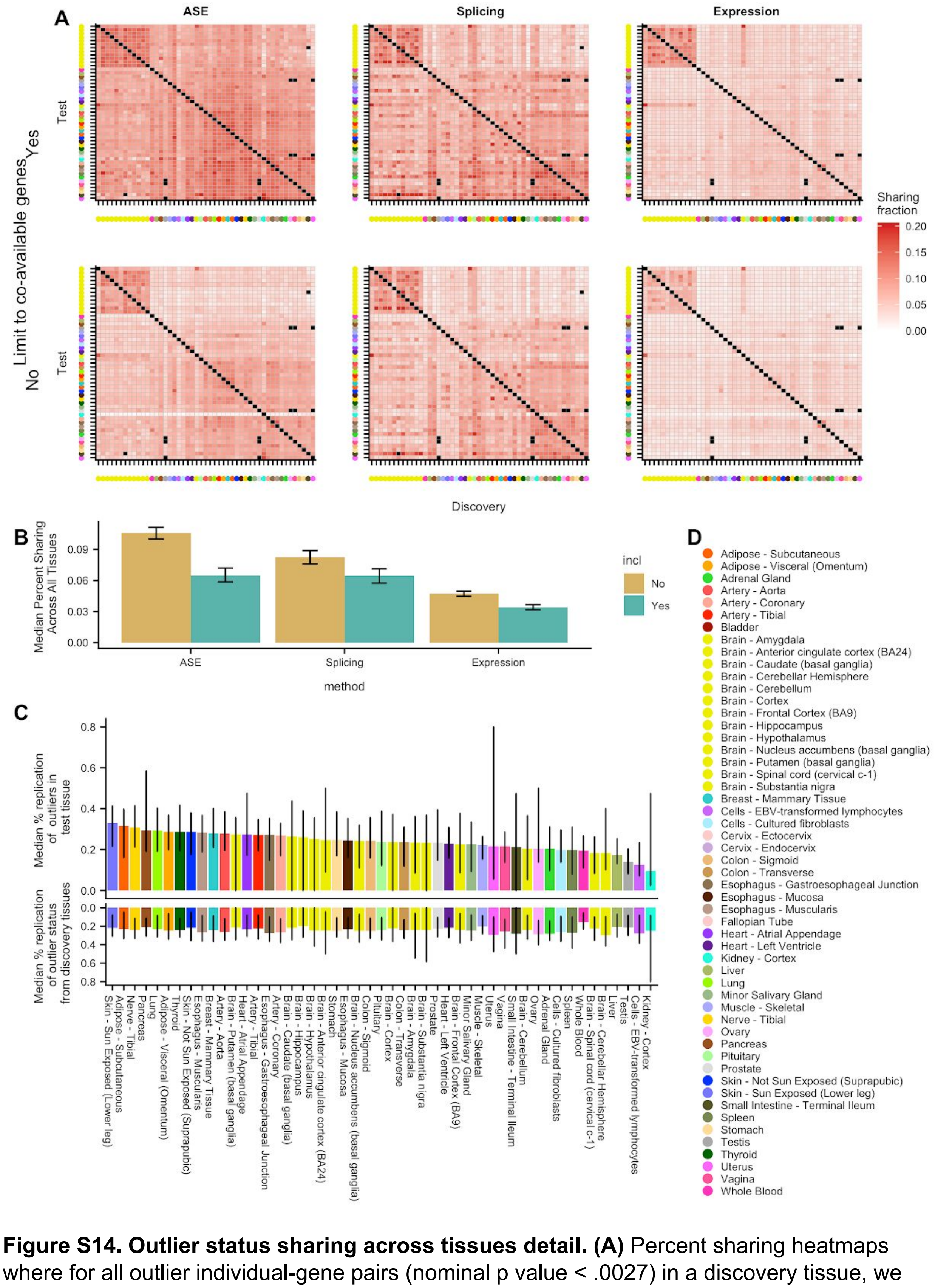


**Figure S14. Outlier status sharing across tissues detail. (A)** Percent sharing heatmaps where for all outlier individual-gene pairs (nominal p value < .0027) in a discovery tissue, we measure the percentage of cases where the same individual-gene pair is also an outlier in a test tissue. In the upper row of heatmaps, we limit the analysis to only the genes tested in both tissues, to answer the biological question of how consistent the outlier status is across tissues that co-express a gene. This is the same figure as in the main text. The lower row of heatmaps considers a missing datapoint as a non-shared outlier status, and addresses the utility of each method in diagnosing expression outlier status in a tissue of interest using a different tissue as a proxy. **(B)** Median percent sharing across all tissue-tissue pairs (± 95% bootstrap confidence interval), with and without considering missing values as "non-shared". aseOutliers are affected the most by missing values. **(C)** Median replication percentage of aseOutlier status in one discovery tissue across all test tissues (top), and median replication percentage of outlier status in one test tissues across all discovery tissues (bottom). The black bars indicate the observed range of values across all individuals. Here, outlier status is declared when a gene has a Benjamini-Hochberg corrected p-value < .05. While for consistency between the three transcriptome outlier methods we use a high significance threshold on the nominal p-values in all other analyses, the FDR correction is the recommended approach when using ANEVA-DOT p-values in most applications. We observe a considerably higher rate of outlier status sharing, when considering genes passing false discovery rate correction. **(D)** The GTEx tissue color key.

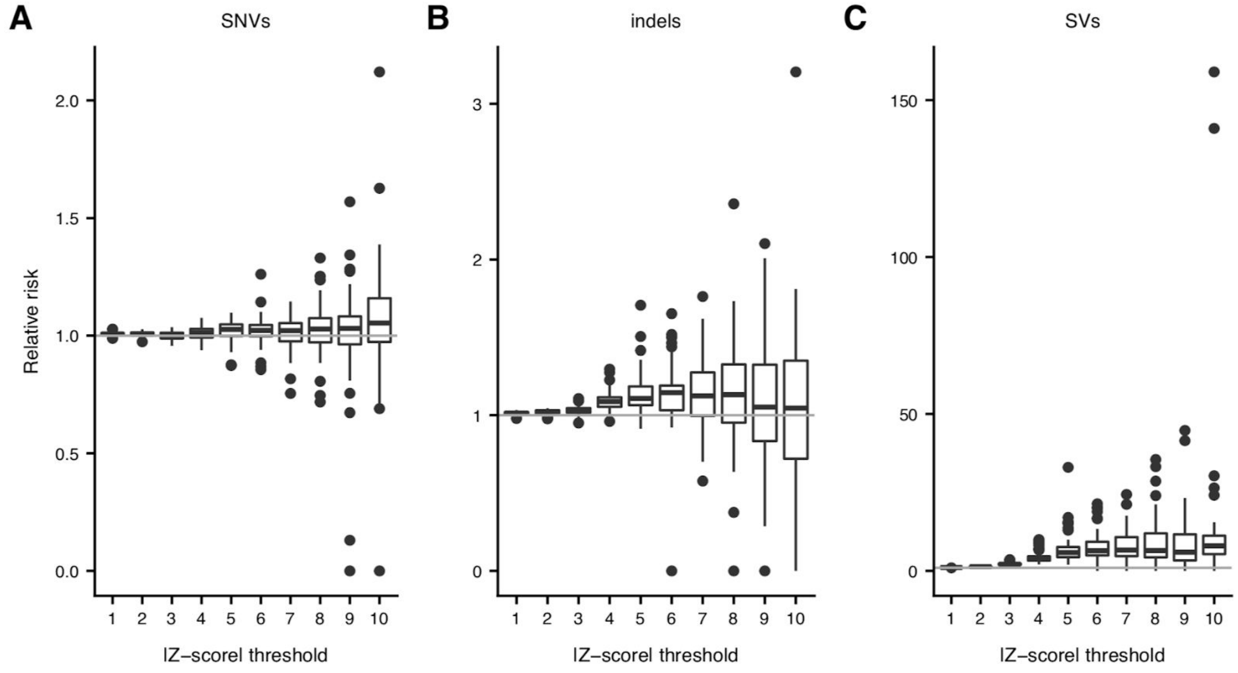


**Figure S15. Single tissue eOutlier enrichments across thresholds.** Relative risk estimates for nearby rare SNVs **(A)**, indels **(B)** and SVs **(C)** in single-tissue outliers vs controls using |Z-score| thresholds between Z=1 and Z=10, with each point representing a single tissue.

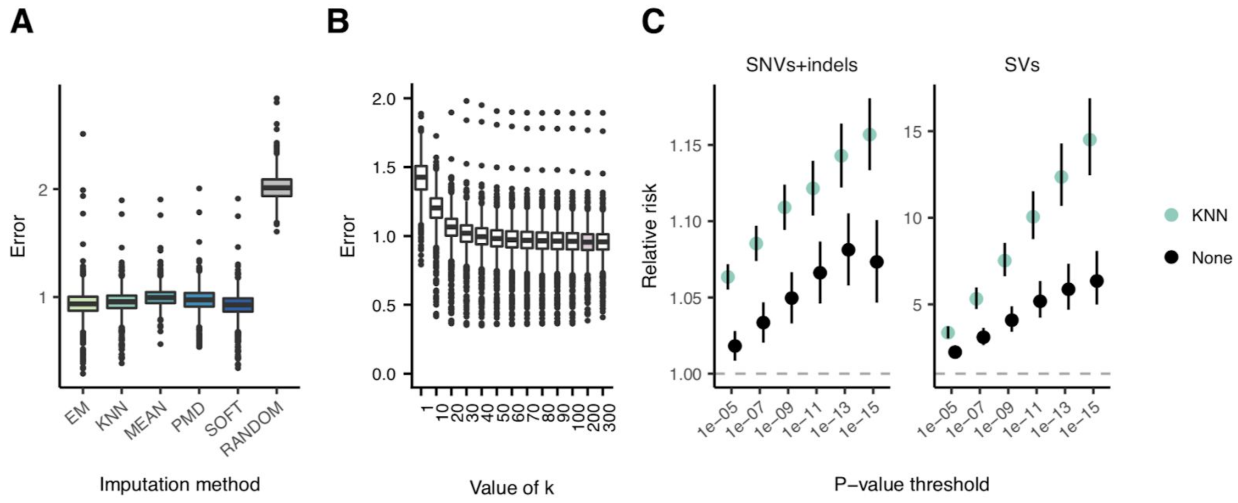


**Figure S16. Comparison of imputation methods and correlation outlier enrichments. (A)** Reconstruction error across genes when holding out 10% of known expression values for various imputation approaches. **(B)** Reconstruction error across genes for different values of k when performing k-nearest neighbors imputation per gene, with the pink box highlighting the value with the lowest error. **(C)** Relative risk of a rare SNV/indel or rare SV nearby correlation outliers called using covariance matrices estimated using KNN-imputed expression data across varying thresholds, as compared to an equal number of outliers called by estimating the covariance matrix from complete entries, without imputation. Many more outliers are identified as compared to the median Z-score approach, particularly at the less stringent thresholds.

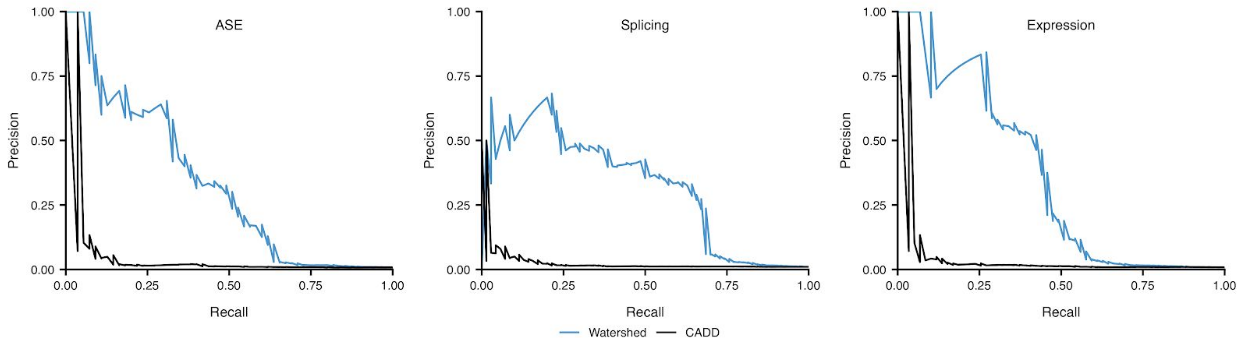


**Figure S17. Precision recall curves for Watershed and CADD.** Precision-recall curves comparing performance of Watershed and CADD (colors) using held out pairs of individuals for all three median outlier signals.

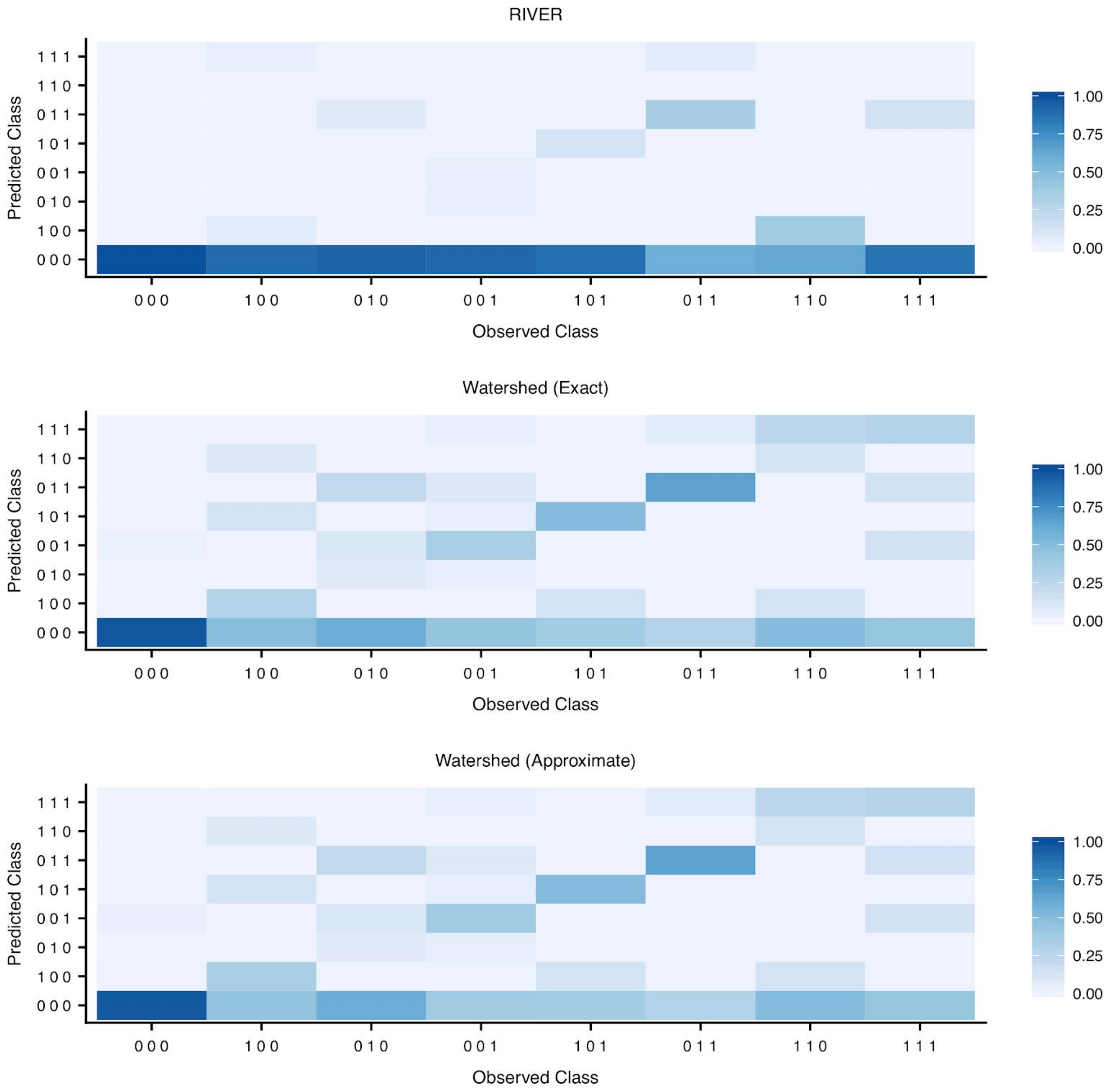


**Figure S18. Watershed confusion matrices.** Confusion matrices comparing performance of RIVER (top), Watershed with parameters optimized via exact inference (middle), and Watershed with parameters optimized via approximate inference (bottom) in jointly predicting outlier status of all three outlier signals (class) using held out pairs of individuals. The first element of the binary class abbreviations represents median splicing outlier status, the second element of the class abbreviations represents median expression outlier status, and the third element of the class abbreviations represents ASE outlier status. An observed class of “1 0 1” therefor corresponds to a sample that is an outlier for splicing and ASE, but not expression. The predicted class of a sample is the class (out of the 8 classes) that has the largest posterior probability. Columns each heatmap are normalized to sum to one.

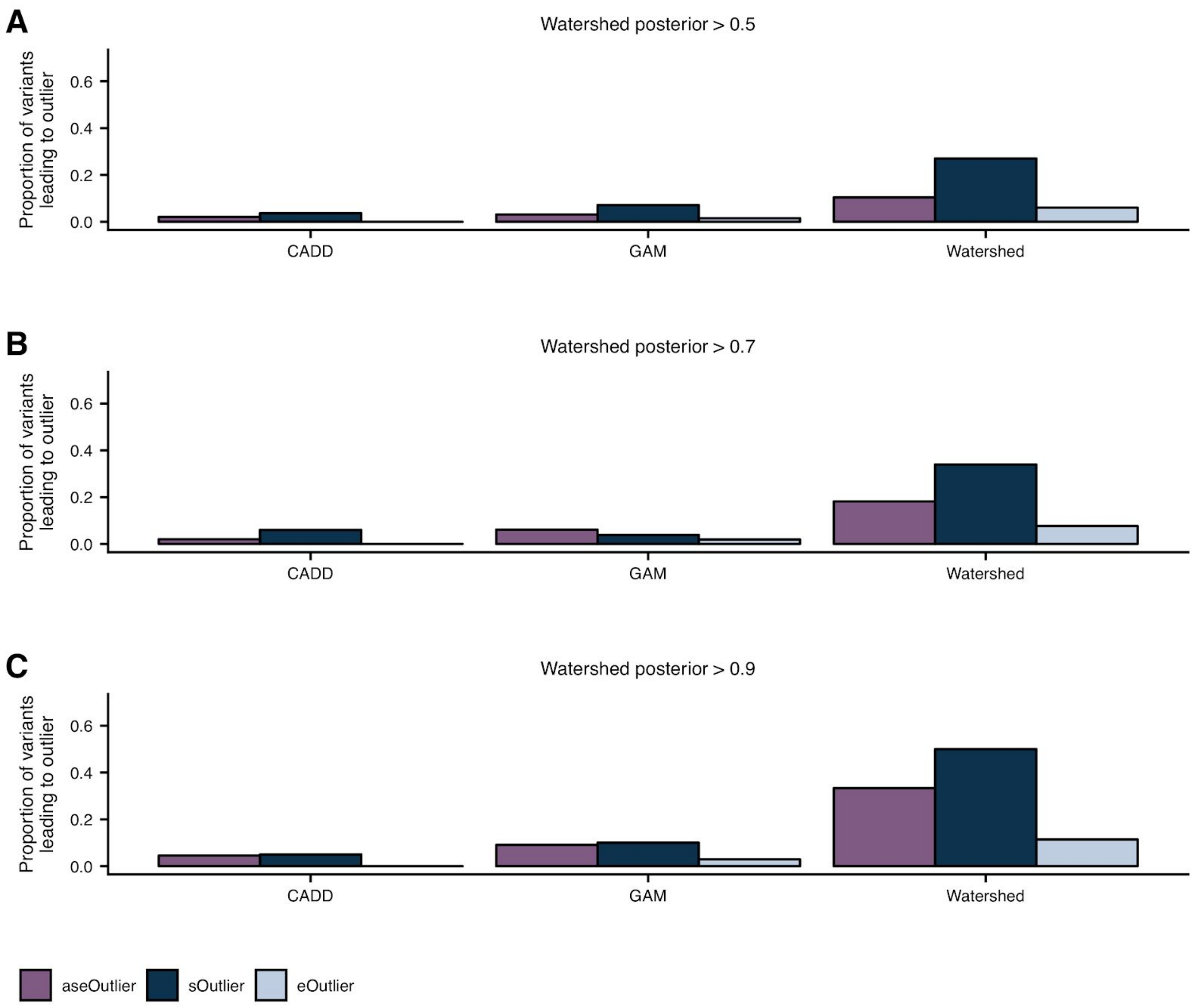


**Figure S19. Prioritization of variants that lead to outliers with Watershed.** The proportion of rare variants, with Watershed posterior probability greater than 0.5 **(A)**, 0.7 **(B)**, 0.9 **(C)** (right), with GAM probability greater than a threshold set to match the number of Watershed variants for each outlier signal (center), and with CADD score greater than a threshold set to match the number of Watershed variants for each outlier signal (left), that lead to an outlier at a median p-value threshold of 0.0027 across three outlier signals (colors). Watershed, GAM, and CADD models evaluated on held-out pairs of individuals.

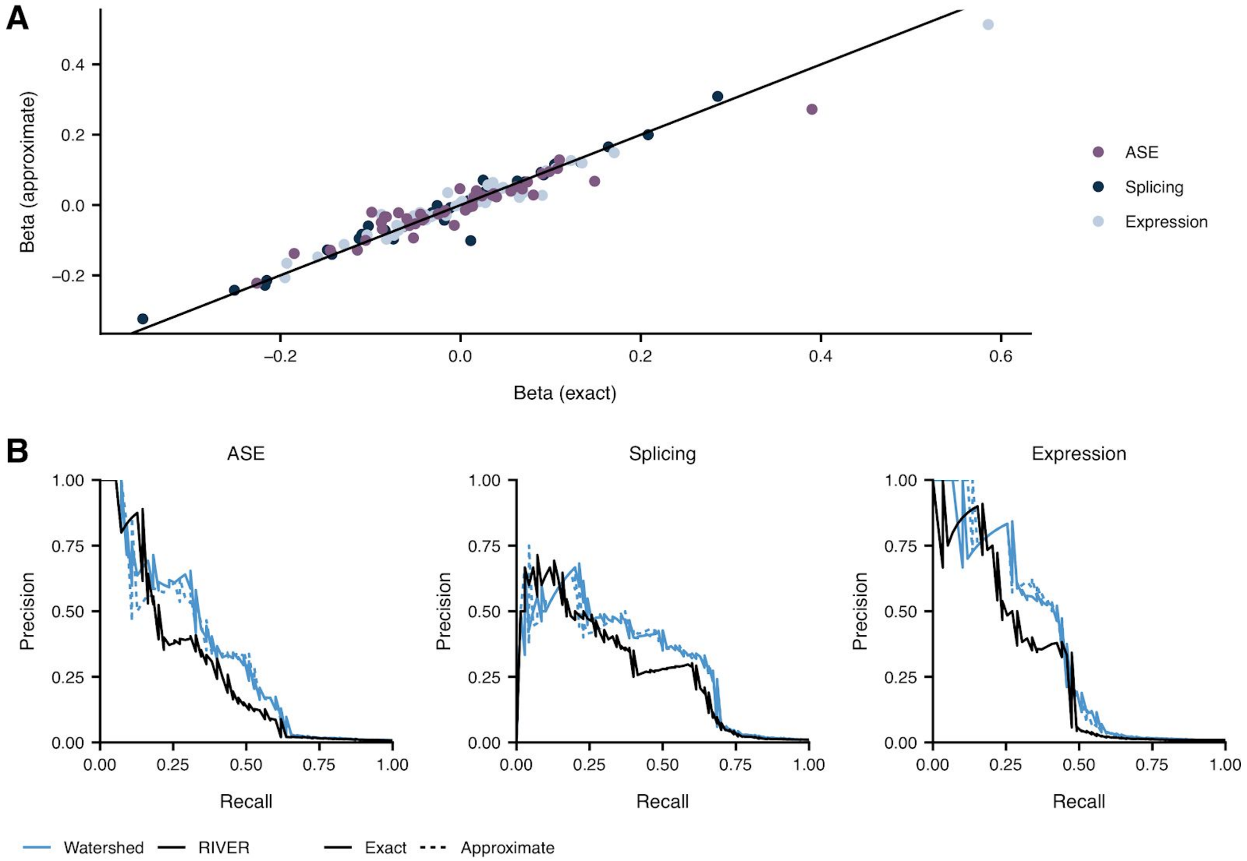


**Figure S20. Comparison of exact and approximate inference in Watershed. (A)** Scatterplot comparing Watershed (applied to median ASE, splicing, and expression outlier signals) genomic annotation coefficients ( $\beta$ ) when model was optimized using exact inference (x-axis) compared to when model was optimized using approximate inference (y-axis) colored by which outlier signal the coefficient predicted. **(B)** Precision-recall curves comparing performance of RIVER, Watershed optimized via exact inference, and Watershed optimized via approximate inference (colors) using held out pairs of individuals for all three median outlier signals.

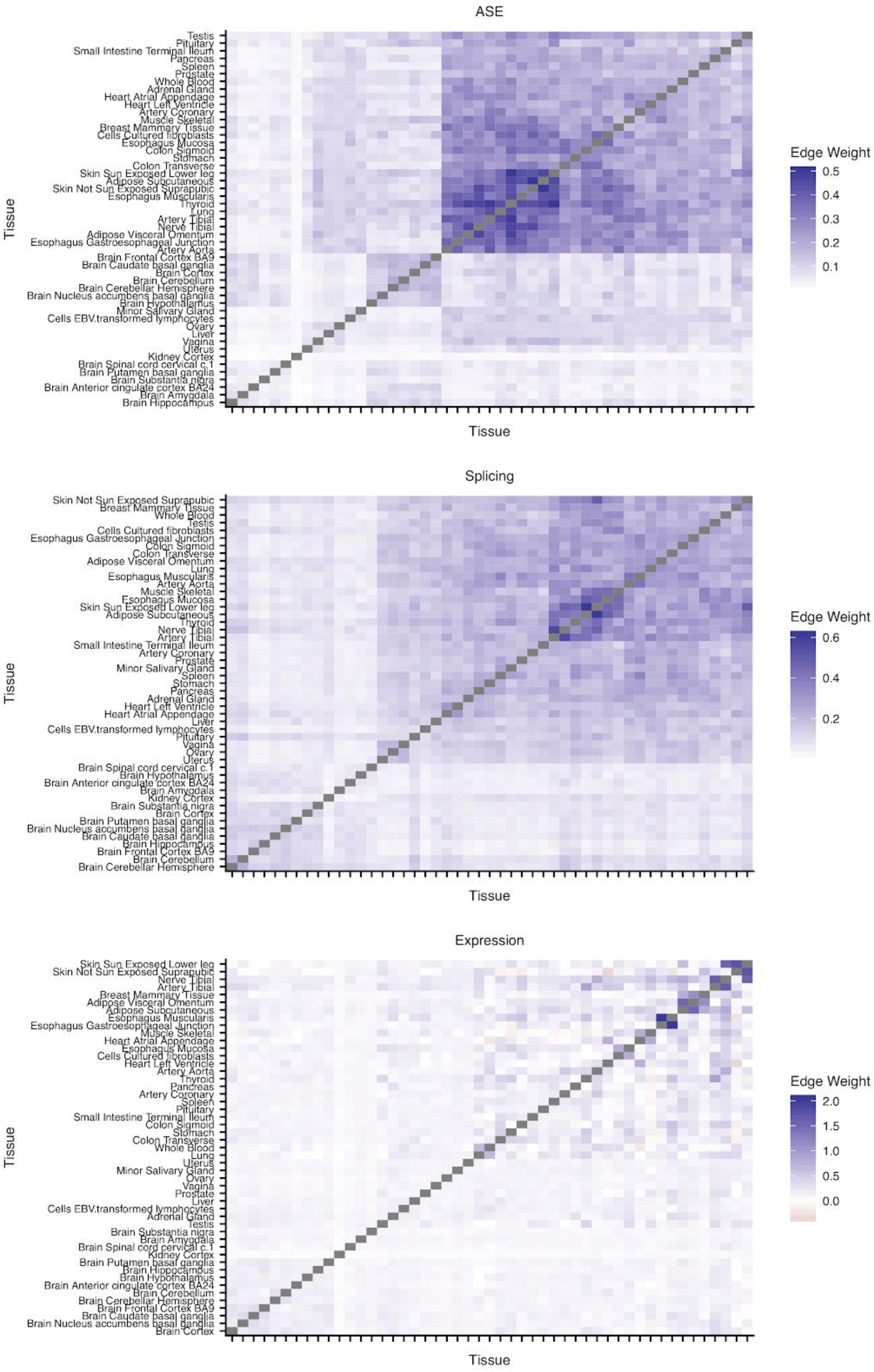
ASE
Testis
Pituitary
Small Intestine Terminal Ileum
Pancreas
Spleen
Prostate
Whole Blood
Adrenal Gland
Heart Atrial Appendage
Heart Left Ventricle
Artery Coronary
Muscle Skeletal
Breast Mammary Tissue
Cells Cultured fibroblasts
Esophagus Mucosa
Colon Sigmoid
Stomach
Colon Transverse
Skin Sun Exposed Lower leg
Adipose Subcutaneous
Skin Not Sun Exposed Suprapubic
Esophagus Muscularis
Thyroid
Lung
Artery Tibial
Nerve Tibial
Adipose Visceral Omentum
Esophagus Gastroesophageal Junction
Artery Aorta
Brain Frontal Cortex BA9
Brain Caudate basal ganglia
Brain Cortex
Brain Cerebellum
Brain Cerebellar Hemisphere
Brain Nucleus accumbens basal ganglia
Brain Hypothalamus
Minor Salivary Gland
Cells EBV.transformed lymphocytes
Ovary
Liver
Vagina
Uterus
Kidney Cortex
Brain Spinal cord cervical c.1
Brain Putamen basal ganglia
Brain Substantia nigra
Brain Anterior cingulate cortex BA24
Brain Amygdala
Brain Hippocampus
Tissue
Tissue
Edge Weight
0.5
0.4
0.3
0.2
0.1
Splicing
Skin Not Sun Exposed Suprapubic
Breast Mammary Tissue
Whole Blood
Testis
Cells Cultured fibroblasts
Esophagus Gastroesophageal Junction
Colon Sigmoid
Colon Transverse
Adipose Visceral Omentum
Lung
Esophagus Muscularis
Artery Aorta
Muscle Skeletal
Esophagus Mucosa
Skin Sun Exposed Lower leg
Adipose Subcutaneous
Thyroid
Nerve Tibial
Artery Tibial
Small Intestine Terminal Ileum
Artery Coronary
Prostate
Minor Salivary Gland
Spleen
Stomach
Pancreas
Adrenal Gland
Heart Left Ventricle
Heart Atrial Appendage
Liver
Cells EBV.transformed lymphocytes
Pituitary
Vagina
Ovary
Uterus
Brain Spinal cord cervical c.1
Brain Hypothalamus
Brain Anterior cingulate cortex BA24
Brain Amygdala
Kidney Cortex
Brain Substantia nigra
Brain Cortex
Brain Putamen basal ganglia
Brain Nucleus accumbens basal ganglia
Brain Caudate basal ganglia
Brain Hippocampus
Brain Frontal Cortex BA9
Brain Cerebellum
Brain Cerebellar Hemisphere
Tissue
Tissue
Edge Weight
0.6
0.4
0.2
Expression
Skin Sun Exposed Lower leg
Skin Not Sun Exposed Suprapubic
Nerve Tibial
Artery Tibial
Breast Mammary Tissue
Adipose Visceral Omentum
Adipose Subcutaneous
Esophagus Muscularis
Esophagus Gastroesophageal Junction
Muscle Skeletal
Heart Atrial Appendage
Esophagus Mucosa
Cells Cultured fibroblasts
Heart Left Ventricle
Artery Aorta
Thyroid
Pancreas
Artery Coronary
Spleen
Pituitary
Small Intestine Terminal Ileum
Colon Sigmoid
Stomach
Colon Transverse
Whole Blood
Lung
Uterus
Minor Salivary Gland
Ovary
Vagina
Prostate
Liver
Cells EBV.transformed lymphocytes
Adrenal Gland
Testis
Brain Substantia nigra
Brain Amygdala
Brain Spinal cord cervical c.1
Kidney Cortex
Brain Putamen basal ganglia
Brain Hippocampus
Brain Hypothalamus
Brain Anterior cingulate cortex BA24
Brain Cerebellum
Brain Cerebellar Hemisphere
Brain Frontal Cortex BA9
Brain Caudate basal ganglia
Brain Nucleus accumbens basal ganglia
Brain Cortex
Tissue
Tissue
Edge Weight
2.0
1.5
1.0
0.5
0.0

**Figure S21. Tissue-specific Watershed edge weights.** Learned tissue-specific Watershed edge weights ( θ ) between pairs of tissue-specific outlier signals after training Watershed on ASE (top), splicing (middle), and expression (bottom) outliers across single tissues.

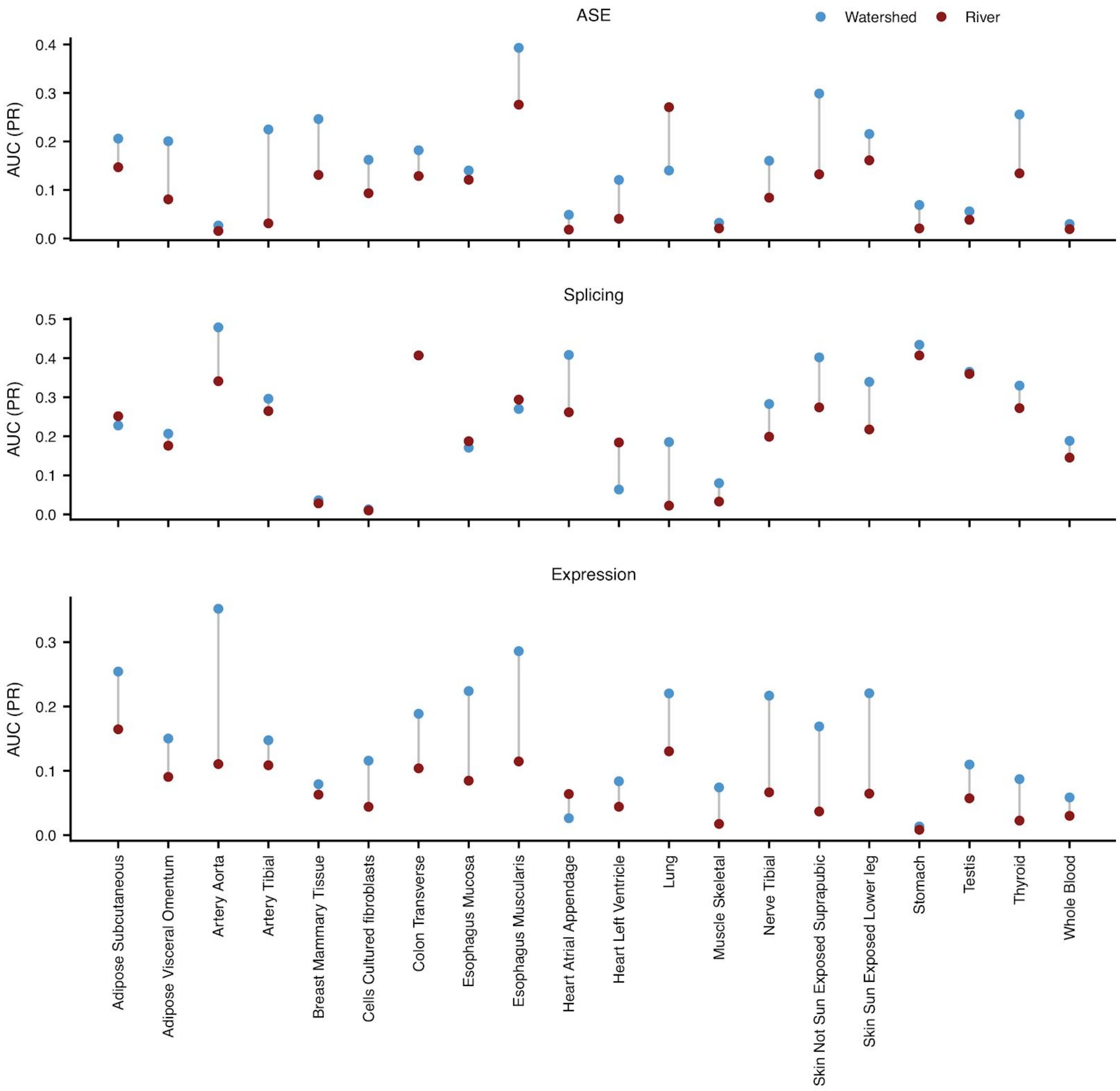


**Figure S22. Area under precision recall curves in single tissues.** Area under precision recall curves (AUC (PR); y-axis) in a single tissue (x-axis) for Watershed (blue) and RIVER (red) when applied outliers across single tissues for all 3 outlier types (rows). Precision recall curves in each tissue generated using held out pairs of individuals where both individuals share the same rare variant and have observed outlier signal for the gene of interest. We limit to tissues that have at least 5 held out pairs of individuals that have outlier labels in ASE, splicing, and expression.

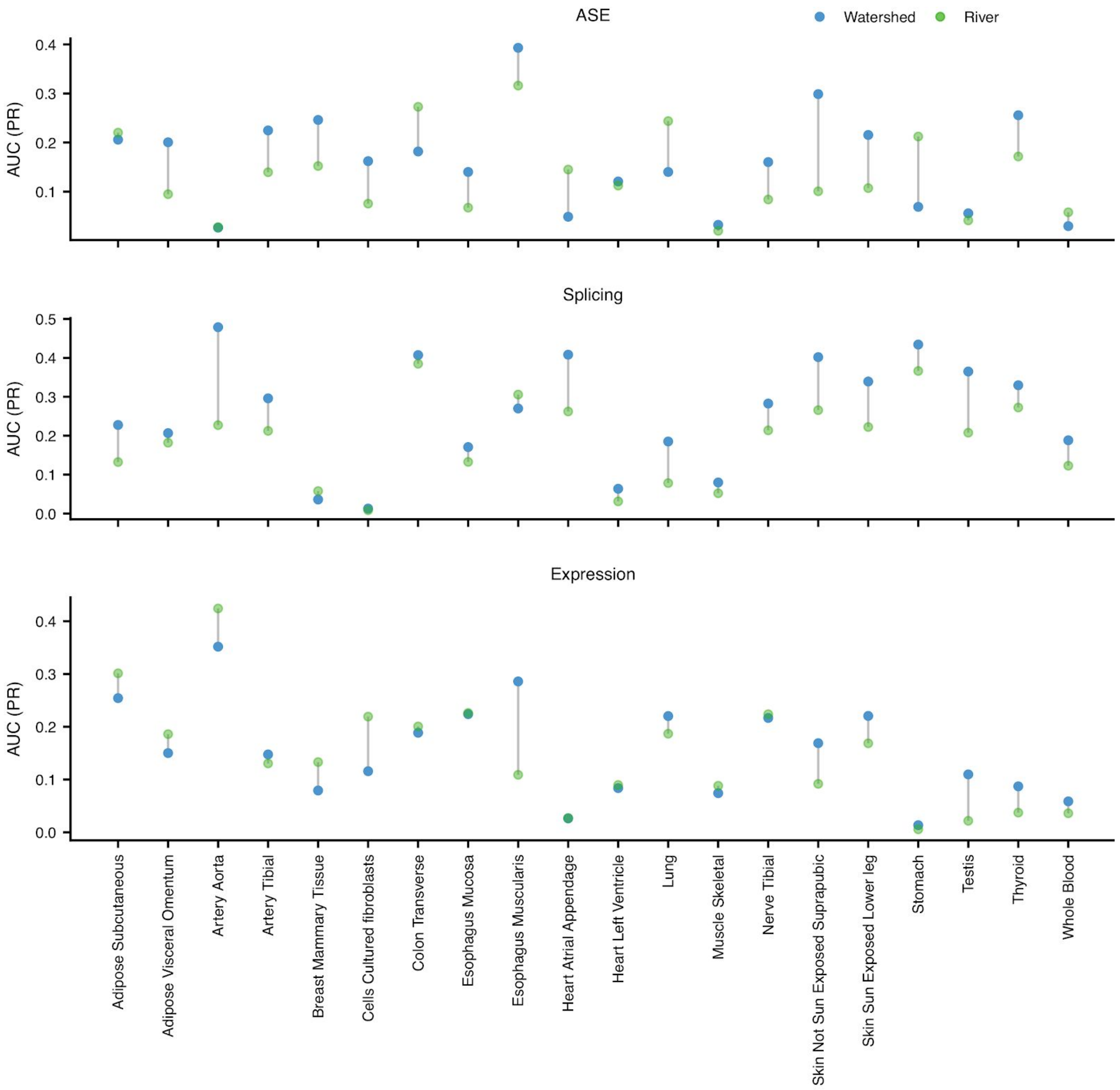


**Figure S23. Area under precision recall curves in single tissues.** Area under precision recall curves evaluated on outlier calls in a single tissue (x-axis) for each of the three outlier types (rows) based on a Watershed model trained across single tissues (blue) and a RIVER model trained on the median outlier signal (green). Precision recall curves in each tissue generated using held out pairs of individuals where both individuals share the same rare variant and have observed outlier signal for the gene of interest. We limit to tissues that have at least 5 held out pairs of individuals that have outlier labels in ASE, splicing, and expression.

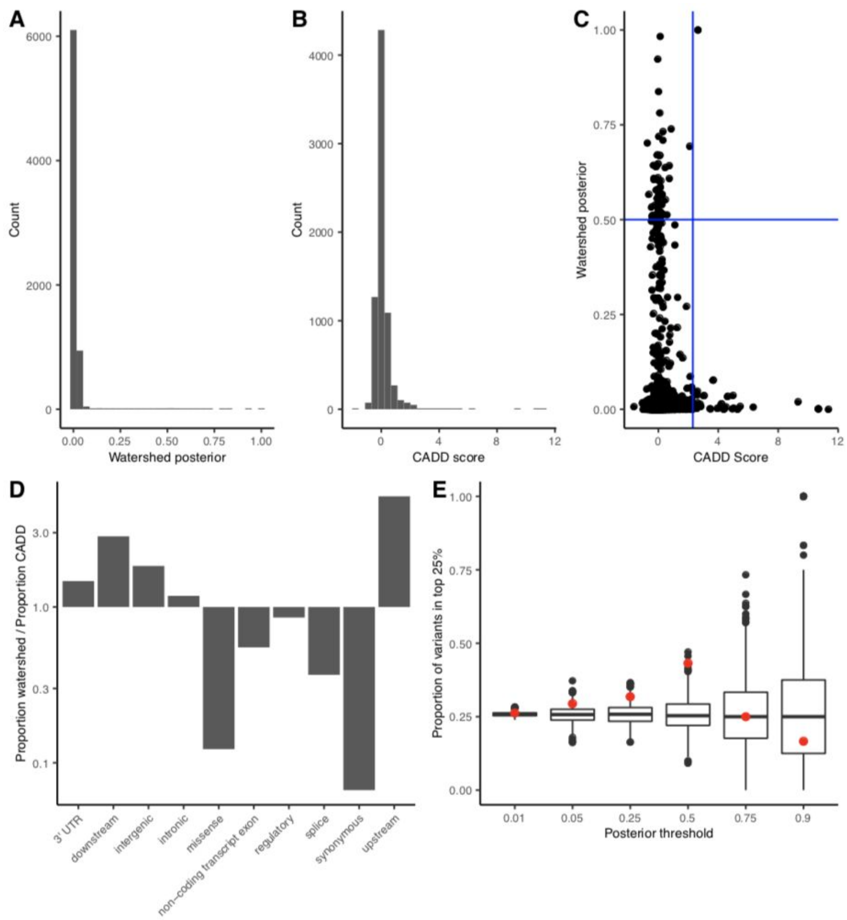


**Figure S24. High CADD and Watershed variants in UKBB. (A)** Distribution of the maximum Watershed posterior per variant for the set of variants in co-localized regions tested by Watershed and in UKBB. **(B)** Distribution of CADD scores per variant for the same set of variants in co-localized regions tested by Watershed and in UKBB. **(C)** The maximum Watershed posterior vs. CADD score for the tested variants in UKBB. The blue lines represent cut-offs of watershed posterior > 0.5, and the matching CADD threshold, 2.3, to obtain the same number of variants. **(D)** Of the high watershed and CADD variants in colocalized regions, the proportion of Watershed variants belonging to a specific category over the proportion of CADD variants in the same category. The y-axis is log-scaled, so bars below 1 indicate the category is more common in high CADD variants, and vice versa. **(E)** Filtering by the CADD score that returns the same number of variants as the Watershed posterior on the x-axis, and returning the proportion that fall in the top 25% of effect sizes across traits in co-localized regions (red), and the proportion obtained by selecting a random set of tested variants equal in size (black).

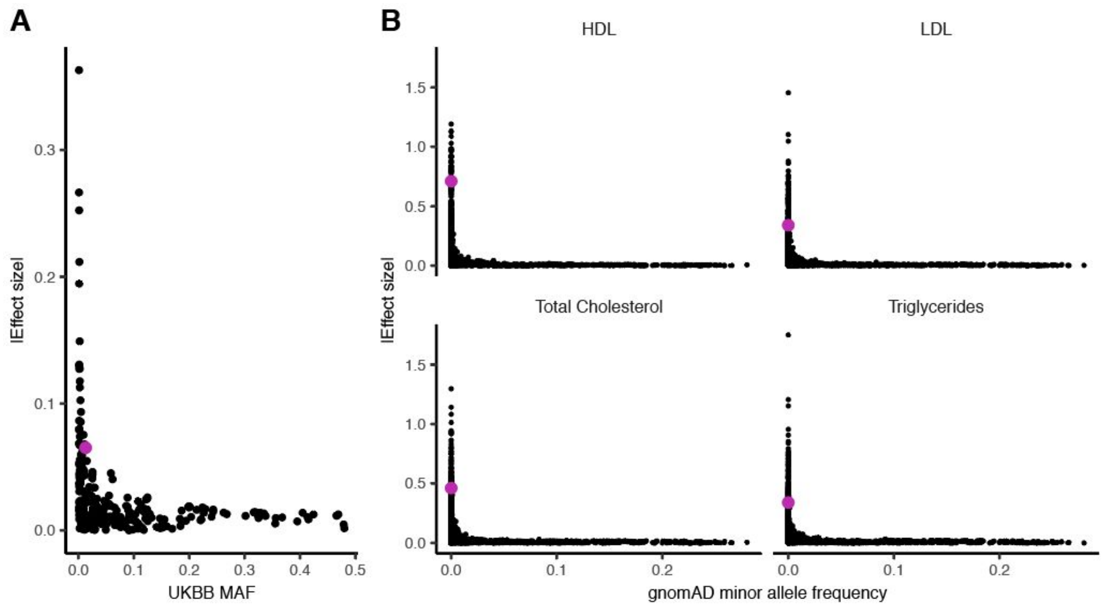


**Figure S25. Distribution of rs564796245 effect sizes in UKBB and MVP. (A)** The UKBB MAF vs. absolute value of the effect size on self-reported high cholesterol for all variants within 10kb of the high Watershed variant, in pink, rs564796245. **(B)** All variants within a 250kb window of the same variant tested for four related traits in the MVP cohort. The variant has a minor allele count of 11 in MVP, and for the set of rare variants tested in this window with a gnomAD non-Finnish European AF < 0.1%, it falls in the 99th percentile for HDL, 95th for LDL, 97th for Total Cholesterol, and 95th for Triglycerides.

#### Supplementary Tables

**Supplementary Table 1. Tissue mapping for Roadmap to GTEx.** Table mapping tissues collected in GTEx to equivalent tissues assayed in the Epigenomics Roadmap project. This includes 12 unique Roadmap tissues and 14 unique GTEx tissues, with some different GTEx tissues mapping to the same Roadmap tissue.

**Supplementary Table 2. UKBB traits and colocalizations.** Table of the 34 UKBB traits included in our analysis and the number of colocalized genes and rare GTEx variants associated with each trait that overlap those tested in the UKBB dataset.

**Supplementary Table 3. High Watershed variants with high effect sizes.** Table of the rare GTEx variants that had both high Watershed scores and high trait effect sizes for the set of UKBB traits tested. This includes the variant, gene, Watershed score, trait, effect size, and the effect size percentile.

**Supplementary Table 4. Watershed genomic annotations.** Table summarizing the 47 genomic annotations used in Watershed. This includes a description of each annotation, the source of each annotation, the imputation value used for each annotation (if the annotation was undefined for a particular variant), and the transformation used to aggregate across all SNVs mapped to (gene, individual) pair for each annotation (only applicable if a (gene, individual) pair had more than one SNV mapped to the gene).

#### GTEx Consortium Information

**Laboratory and Data Analysis Coordinating Center (LDACC):** François Aguet[1], Shankara Anand[1], Kristin G Ardlie[1], Stacey Gabriel[1], Gad Getz[1,2], Aaron Graubert[1], Kane Hadley[1], Robert E Handsaker[3,4,5], Katherine H Huang[1], Seva Kashin[3,4,5], Xiao Li[1], Daniel G MacArthur[4,6], Samuel R Meier[1], Jared L Nedzel[1], Duyen Y Nguyen[1], Ayellet V Segrè[1,7], Ellen Todres[1]
**Analysis Working Group (funded by GTEx project grants):** François Aguet[1], Shankara Anand[1], Kristin G Ardlie[1], Brunilda Balliu[8], Alvaro N Barbeira[9], Alexis Battle[10,11], Rodrigo Bonazzola[9], Andrew Brown[12,13], Christopher D Brown[14], Stephane E Castel[15,16], Don Conrad[17,18], Daniel J Cotter[19], Nancy Cox[20], Sayantan Das[21], Olivia M de Goede[19], Emmanouil T Dermitzakis[22,23,24], Barbara E Engelhardt[25,26], Eleazar Eskin[27], Tiffany Y Eulalio[28], Nicole M Ferraro[28], Elise Flynn[15,16], Laure Fresard[29], Eric R Gamazon[30,31,32,20], Diego Garrido-Martín[33], Nicole R Gay[19], Gad Getz[1,2], Aaron Graubert[1], Roderic Guigó[33,34], Kane Hadley[1], Andrew R Hamel[7,1], Robert E Handsaker[3,4,5], Yuan He[10], Paul J Hoffman[15], Farhad Hormozdiari[35,1], Lei Hou[36,1], Katherine H Huang[1], Hae Kyung Im[9], Brian Jo[25,26], Silva Kasela[15,16], Seva Kashin[3,4,5], Manolis Kellis[36,1], Sarah Kim-Hellmuth[15,16,37], Alan Kwong[21], Tuuli Lappalainen[15,16], Xiao Li[1], Xin Li[29], Yanyu Liang[9], Daniel G MacArthur[4,6], Serghei Mangul[27,38], Samuel R Meier[1], Pejman Mohammadi[15,16,39,40], Stephen B Montgomery[19,29], Manuel Muñoz-Aguirre[33,41], Daniel C Nachun[29], Jared L Nedzel[1], Duyen Y Nguyen[1], Andrew B Nobel[42], Meritxell Oliva[9,43], YoSon Park[14,44], Yongjin Park[36,1], Princy Parsana[11], Ferran Reverter[45], John M Rouhana[7,1], Chiara Sabatti[46], Ashis Saha[11], Ayellet V Segrè[1,7], Andrew D Skol[9,47], Matthew Stephens[48], Barbara E Stranger[9,49], Benjamin J Strober[10], Nicole A Teran[29], Ellen Todres[1], Ana Viñuela[50,22,23,24], Gao Wang[48], Xiaoquan Wen[21], Fred Wright[51], Valentin Wucher[33], Yuxin Zou[52]
**Analysis Working Group (not funded by GTEx project grants):** Pedro G Ferreira[53,54,55], Gen Li[56], Marta Melé[57], Esti Yeger-Lotem[58,59]
**Leidos Biomedical - Project Management:** Mary E Barcus[60], Debra Bradbury[61], Tanya Krubit[61], Jeffrey A McLean[61], Liqun Qi[61], Karna Robinson[61], Nancy V Roche[61], Anna M Smith[61], Leslie Sobin[61], David E Tabor[61], Anita Undale[61]
**Biospecimen collection source sites:** Jason Bridge[62], Lori E Brigham[63], Barbara A Foster[64], Bryan M Gillard[64], Richard Hasz[65], Marcus Hunter[66], Christopher Johns[67], Mark Johnson[68], Ellen Karasik[64], Gene Kopen[69], William F Leinweber[69], Alisa McDonald[69], Michael T Moser[64], Kevin Myer[66], Kimberley D Ramsey[64], Brian Roe[66], Saboor Shad[69], Jeffrey A Thomas[69,68], Gary Walters[68], Michael Washington[68], Joseph Wheeler[67]
**Biospecimen core resource:** Scott D Jewell[70], Daniel C Rohrer[70], Dana R Valley[70]
**Brain bank repository:** David A Davis[71], Deborah C Mash[71]
**Pathology:** Mary E Barcus[60], Philip A Branton[72], Leslie Sobin[61]
**ELSI study:** Laura K Barker[73], Heather M Gardiner[73], Maghboeba Mosavel[74], Laura A Siminoff[73]
**Genome Browser Data Integration & Visualization:** Paul Flicek[75], Maximilian Haeussler[76], Thomas Juettemann[75], W James Kent[76], Christopher M Lee[76], Conner C Powell[76], Kate R Rosenbloom[76], Magali Ruffier[75], Dan Sheppard[75], Kieron Taylor[75], Stephen J Trevanion[75], Daniel R Zerbino[75]
**eGTEx groups:** Nathan S Abell[19], Joshua Akey[77], Lin Chen[43], Kathryn Demanelis[43], Jennifer A Doherty[78], Andrew P Feinberg[79], Kasper D Hansen[80], Peter F Hickey[81], Lei Hou[36,1], Farzana Jasmine[43], Lihua Jiang[19], Rajinder Kaul[82,83], Manolis Kellis[36,1], Muhammad G Kibriya[43], Jin Billy

Li[19], Qin Li[19], Shin Lin[84], Sandra E Linder[19], Stephen B Montgomery[29,19], Meritxell Oliva[9,43], Yongjin Park[36,1], Brandon L Pierce[43], Lindsay F Rizzardi[85], Andrew D Skol[9,47], Kevin S Smith[29], Michael Snyder[19], John Stamatoyannopoulos[82,86], Barbara E Stranger[9,49], Hua Tang[19], Meng Wang[19]
**NIH program management:** Philip A Branton[72], Latarsha J Carithers[72,87], Ping Guan[72], Susan E Koester[88], A. Roger Little[89], Helen M Moore[72], Concepcion R Nierras[90], Abhi K Rao[72], Jimmie B Vaught[72], Simona Volpi[91]
**Affiliations**
1. The Broad Institute of MIT and Harvard, Cambridge, MA, USA
2. Cancer Center and Department of Pathology, Massachusetts General Hospital, Boston, MA, USA
3. Department of Genetics, Harvard Medical School, Boston, MA, USA
4. Program in Medical and Population Genetics, The Broad Institute of Massachusetts Institute of Technology and Harvard University, Cambridge, MA, USA
5. Stanley Center for Psychiatric Research, Broad Institute, Cambridge, MA, USA
6. Analytic and Translational Genetics Unit, Massachusetts General Hospital, Boston, MA, USA
7. Ocular Genomics Institute, Massachusetts Eye and Ear, Harvard Medical School, Boston, MA, USA
8. Department of Biomathematics, University of California, Los Angeles, Los Angeles, CA, USA
9. Section of Genetic Medicine, Department of Medicine, The University of Chicago, Chicago, IL, USA
10. Department of Biomedical Engineering, Johns Hopkins University, Baltimore, MD, USA
11. Department of Computer Science, Johns Hopkins University, Baltimore, MD, USA
12. Department of Genetic Medicine and Development, University of Geneva Medical School, Geneva, Switzerland
13. Population Health and Genomics, University of Dundee, Dundee, Scotland, UK
14. Department of Genetics, University of Pennsylvania, Perelman School of Medicine, Philadelphia, PA, USA
15. New York Genome Center, New York, NY, USA
16. Department of Systems Biology, Columbia University, New York, NY, USA
17. Department of Genetics, Washington University School of Medicine, St. Louis, Missouri, USA
18. Department of Pathology & Immunology, Washington University School of Medicine, St. Louis, Missouri, USA
19. Department of Genetics, Stanford University, Stanford, CA, USA
20. Division of Genetic Medicine, Department of Medicine, Vanderbilt University Medical Center, Nashville, TN, USA
21. Department of Biostatistics, University of Michigan, Ann Arbor, MI, USA
22. Department of Genetic Medicine and Development, University of Geneva Medical School, Geneva, Switzerland
23. Institute for Genetics and Genomics in Geneva (iGE3), University of Geneva, Geneva, Switzerland
24. Swiss Institute of Bioinformatics, Geneva, Switzerland
25. Department of Computer Science, Princeton University, Princeton, NJ, USA
26. Center for Statistics and Machine Learning, Princeton University, Princeton, NJ, USA
27. Department of Computer Science, University of California, Los Angeles, Los Angeles, CA, USA

28. Program in Biomedical Informatics, Stanford University School of Medicine, Stanford, CA, USA
29. Department of Pathology, Stanford University, Stanford, CA, USA
30. Data Science Institute, Vanderbilt University, Nashville, TN, USA
31. Clare Hall, University of Cambridge, Cambridge, UK
32. MRC Epidemiology Unit, University of Cambridge, Cambridge, UK
33. Centre for Genomic Regulation (CRG), The Barcelona Institute for Science and Technology, Barcelona, Catalonia, Spain
34. Universitat Pompeu Fabra (UPF), Barcelona, Catalonia, Spain
35. Department of Epidemiology, Harvard T.H. Chan School of Public Health, Boston, MA, USA
36. Computer Science and Artificial Intelligence Laboratory, Massachusetts Institute of Technology, Cambridge, MA, USA
37. Statistical Genetics, Max Planck Institute of Psychiatry, Munich, Germany
38. Department of Clinical Pharmacy, School of Pharmacy, University of Southern California, Los Angeles, CA, USA
39. Scripps Research Translational Institute, La Jolla, CA, USA
40. Department of Integrative Structural and Computational Biology, The Scripps Research Institute, La Jolla, CA, USA
41. Department of Statistics and Operations Research, Universitat Politècnica de Catalunya (UPC), Barcelona, Catalonia, Spain
42. Department of Statistics and Operations Research and Department of Biostatistics, University of North Carolina, Chapel Hill, NC, USA
43. Department of Public Health Sciences, The University of Chicago, Chicago, IL, USA
44. Department of Systems Pharmacology and Translational Therapeutics, University of Pennsylvania, Perelman School of Medicine, Philadelphia, PA, USA
45. Department of Genetics, Microbiology and Statistics, University of Barcelona, Barcelona. Spain.
46. Departments of Biomedical Data Science and Statistics, Stanford University, Stanford, CA, USA
47. Department of Pathology and Laboratory Medicine, Ann & Robert H. Lurie Children's Hospital of Chicago, Chicago, IL, USA
48. Department of Human Genetics, University of Chicago, Chicago, IL, USA
49. Center for Genetic Medicine, Department of Pharmacology, Northwestern University, Feinberg School of Medicine, Chicago, IL, USA
50. Department of Twin Research and Genetic Epidemiology, King’s College London, London, UK
51. Bioinformatics Research Center and Departments of Statistics and Biological Sciences, North Carolina State University, Raleigh, NC, USA
52. Department of Statistics, University of Chicago, Chicago, IL, USA
53. Department of Computer Sciences, Faculty of Sciences, University of Porto, Porto, Portugal
54. Instituto de Investigação e Inovação em Saúde, Universidade do Porto, Porto, Portugal
55. Institute of Molecular Pathology and Immunology, University of Porto, Porto, Portugal
56. Columbia University Mailman School of Public Health, New York, NY, USA
57. Life Sciences Department, Barcelona Supercomputing Center, Barcelona, Spain
58. Department of Clinical Biochemistry and Pharmacology, Ben-Gurion University of the Negev, Beer-Sheva, Israel
59. National Institute for Biotechnology in the Negev, Beer-Sheva, Israel
60. Leidos Biomedical, Frederick, MD, USA

61. Leidos Biomedical, Rockville, MD, USA
62. UNYTS, Buffalo, NY, USA
63. Washington Regional Transplant Community, Annandale, VA, USA
64. Therapeutics, Roswell Park Comprehensive Cancer Center, Buffalo, NY, USA
65. Gift of Life Donor Program, Philadelphia, PA, USA
66. LifeGift, Houston, TX, USA
67. Center for Organ Recovery and Education, Pittsburgh, PA, USA
68. LifeNet Health, Virginia Beach, VA. USA
69. National Disease Research Interchange, Philadelphia, PA, USA
70. Van Andel Research Institute, Grand Rapids, MI, USA
71. Department of Neurology, University of Miami Miller School of Medicine, Miami, FL, USA
72. Biorepositories and Biospecimen Research Branch, Division of Cancer Treatment and Diagnosis, National Cancer Institute, Bethesda, MD, USA
73. Temple University, Philadelphia, PA, USA
74. Virgina Commonwealth University, Richmond, VA, USA
75. European Molecular Biology Laboratory, European Bioinformatics Institute, Hinxton, United Kingdom
76. Genomics Institute, UC Santa Cruz, Santa Cruz, CA, USA
77. Carl Icahn Laboratory, Princeton University, Princeton, NJ, USA
78. Department of Population Health Sciences, The University of Utah, Salt Lake City, Utah, USA
79. Schools of Medicine, Engineering, and Public Health, Johns Hopkins University, Baltimore, MD, USA
80. Department of Biostatistics, Bloomberg School of Public Health, Johns Hopkins University, Baltimore, MD, USA
81. Department of Medical Biology, The Walter and Eliza Hall Institute of Medical Research, Parkville, Victoria, Australia
82. Altius Institute for Biomedical Sciences, Seattle, WA, USA
83. Division of Genetics, University of Washington, Seattle, WA, University of Washington, Seattle, WA, USA
84. Department of Cardiology, University of Washington, Seattle, WA, USA
85. HudsonAlpha Institute for Biotechnology, Huntsville, AL, USA
86. Genome Sciences, University of Washington, Seattle, WA, USA
87. National Institute of Dental and Craniofacial Research, Bethesda, MD, USA
88. Division of Neuroscience and Basic Behavioral Science, National Institute of Mental Health, National Institutes of Health, Bethesda, MD, USA
89. National Institute on Drug Abuse, Bethesda, MD, USA
90. Office of Strategic Coordination, Division of Program Coordination, Planning and Strategic Initiatives, Office of the Director, National Institutes of Health, Rockville, MD, USA
91. Division of Genomic Medicine, National Human Genome Research Institute, Bethesda, MD, USA

**Funding**

The consortium was funded by GTEx program grants: HHSN268201000029C (F.A., K.G.A., A.V.S., X.Li., E.T., S.G., A.G., S.A., K.H.H., D.Y.N., K.H., S.R.M., J.L.N.), 5U41HG009494 (F.A., K.G.A.), 10XS170 (Subcontract to Leidos Biomedical) (W.F.L., J.A.T., G.K., A.M., S.S., R.H., G.Wa., M.J., M.Wa., L.E.B., C.J., J.W., B.R., M.Hu., K.M., L.A.S., H.M.G., M.Mo., L.K.B.), 10XS171 (Subcontract to Leidos Biomedical) (B.A.F., M.T.M., E.K., B.M.G., K.D.R., J.B.), 10ST1035 (Subcontract to Leidos Biomedical) (S.D.J., D.C.R., D.R.V.), R01DA006227-17

(D.C.M., D.A.D.), Supplement to University of Miami grant DA006227. (D.C.M., D.A.D.), HHSN261200800001E (A.M.S., D.E.T., N.V.R., J.A.M., L.S., M.E.B., L.Q., T.K., D.B., K.R., A.U.), R01MH101814 (M.M-A., V.W., S.B.M., R.G., E.T.D., D.G-M., A.V.), U01HG007593 (S.B.M.), R01MH101822 (C.D.B.), U01HG007598 (M.O., B.E.S.).

**COI**

F.A. is an inventor on a patent application related to TensorQTL; S.E.C. is a co-founder, chief technology officer and stock owner at Variant Bio; E.R.G. is on the Editorial Board of Circulation Research, and does consulting for the City of Hope / Beckman Research Institut; E.T.D. is chairman and member of the board of Hybridstat LTD.; B.E.E. is on the scientific advisory boards of Celsius Therapeutics and Freenome; G.G. receives research funds from IBM and Pharmacyclics, and is an inventor on patent applications related to MuTect, ABSOLUTE, MutSig, POLYSOLVER and TensorQTL; S.B.M. is on the scientific advisory board of Prime Genomics Inc.; D.G.M. is a co-founder with equity in Goldfinch Bio, and has received research support from AbbVie, Astellas, Biogen, BioMarin, Eisai, Merck, Pfizer, and Sanofi-Genzyme; H.K.I. has received speaker honoraria from GSK and AbbVie.; T.L. is a scientific advisory board member of Variant Bio with equity and Goldfinch Bio. P.F. is member of the scientific advisory boards of Fabric Genomics, Inc., and Eagle Genomes, Ltd. P.G.F. is a partner of Bioinf2Bio.
